# Postzygotic and germinal *de novo* mutations in ASD: exploring their biological role

**DOI:** 10.1101/2020.05.21.107987

**Authors:** A Alonso-González, M Calaza, J Amigo, J González-Peñas, Martínez-Regueiro, M Fernández-Prieto, M Parellada, C Rodriguez-Fontenla, A Carracedo

## Abstract

De novo mutations (DNMs), including germinal and postzygotic mutations (PZMs), are a strong source of causality for Autism Spectrum Disorder (ASD). However, the biological processes involved behind them remain unexplored. Our aim was to detect DNMs (germinal and PZMs) in a Spanish ASD cohort (360 trios) and to explore their role across different biological hierarchies (gene, biological pathway, cell and brain areas) using bioinformatic approaches. For the majority of the analysis, a combined cohort (N=2171 trios) with ASC (Autism Sequencing Consortium) previously published data was created. New plausible candidate genes for ASD such as *FMR1* and *NFIA* were found. In addition, genes harboring PZMs were significantly enriched for miR-137 targets in comparison with germinal DNMs that were enriched in GO terms related to synaptic transmission. The expression pattern of genes with PZMs was restricted to early mid-fetal cortex. In contrast, the analysis of genes with germinal DNMs revealed a spatio-temporal window from early to mid-fetal development stages, with expression in the amygdala, cerebellum, cortex and striatum. These results provide evidence of the pathogenic role of PZMs and suggest the existence of distinct mechanisms between PZMs and germinal DNMs that are influencing ASD risk.

## BACKGROUND

Autism Spectrum Disorder (ASD) is a neurodevelopmental disorder (NDD) characterized by deficits in communication and social interaction together with restricted interests and repetitive behaviors^1^. ASD prevalence among children in the United States stands at around 1.5 % and has rapidly risen in recent years. In addition to the core symptoms of ASD, other conditions such as epilepsy or intellectual disability (ID) are often present. Comorbidity is a characteristic of ASD that can appear at any time during child’s development. Since many ASC cases with comorbidity have a clear genetic background and early detection is key for intervention, the genetic diagnosis in this type of cases is a challenge ^2^.

Twin and family studies have estimated ASD heritability to be about 80% and subsequent genetic studies have demonstrated that the largest part of this heritability (50%) is explained by common variation^3,4^. However, *de novo* rare genetic variation (minor allele frequency < 0.1%), including small insertions and deletions (indels), copy number variants and single nucleotide variants confers higher individual risk^5–7^. Germinal *de novo* mutations (DNMs) occur within germ cells and they are transmitted to the offspring when the zygote is formed after fertilization. Thus, every single cell line of the resulting embryo will carry an identical genetic load. Another type of DNMs, postzygotic mutations (PZMs), arise during zygote mitosis, leading to a mosaic of genetically different cell lines^8^. The frequency of mutagenesis and the generation of PZMs is increased prior to gastrulation and neurogenesis^9^.

PZMs involved in ASD pathogenesis are usually detectable through deep sequencing of brain tissues. However, this technique often entails a huge challenge due to the inability to obtain ASD brain samples^10^. By contrast, next generation sequencing technologies are useful to detect mosaic mutations in peripheral blood of affected individuals due to the increase in the depth of coverage^11^. Thus, it is possible to obtain enough sequencing reads containing the reference and the alternate allele to accurately calculate the alternate allele frequency (AAF)^12^. In PZMs, the AAF value shifts from the expected 50/50 ratio for heterozygous germinal mutations. High coverage WES (whole exome sequencing) (depth > 200X) provides enough sensitivity to detect PZMs presenting AAF values as lower as 15%^13,14^. It is worth to note that most WES studies have missed PZMs due to the commonly employed pipelines. The development of new variant calling pipelines is therefore needed and some efforts have been done at this regard.^15–18^.

It has been estimated that 7.5% of DNMs are PZMs that contribute about 4% to the overall architecture of ASD. PZMs have been identified in high-confidence ASD risk genes. Other novel ASD candidate genes such as *KLF16* and *MSANTD2*, were discovered after studying the contribution of PZMs to ASD risk in large collections of ASD probands^18^. This points to the fact that some genes tend to carry a larger number of mutations in a mosaic state than other genes. In addition, a detailed analysis of non-synonymous PZMs has revealed that these variants are mainly found in brain-expressed genes and in Loss-of-function (LoF)-constrained exon. The spatio-temporal analysis across different developmental stages also points to brain areas, like the amygdala, that have not been previously highlighted by other WES studies in which PZMs were not considered^16,18^.

The relevance of PZMs in the pathogenesis of ASD and the biological processes in which genes carrying PZMs are involved, remain largely unexplored. Moreover, the contribution of PZMs to the phenotypic presentation is another subject that should be studied in more detail using large-scale studies. Hence, it is suspected that ASD probands carrying mosaic mutations might be less affected than probands carrying germinal mutations as it happens in other neurodevelopmental disorders such as Proteus syndrome or several brain malformations^19,20^. Therefore, the main aim of this study was to accurately detect DNMs (germinal and PZMs) in a cohort of Spanish trios with ASD (360). The novel DNMs detected in the Spanish cohort were combined with a list of DNMs previously published by the Autism Sequencing Consortium (ASC)^18^ in a cohort of 5947 families (4,032 ASD trios and 1,918 quads) in order to study if different ASD risk genes tend to accumulate one or another type of mutations using different bioinformatic approaches. In addition, the different biological implications of germinal and PZMs in ASD were explored through enrichment analysis approaches, which have not been applied before to this class of mutations across different hierarchical levels (gene, GO terms, neuronal cell types and brain areas) **(Additional file 4. Figure S1)**.

## METHODS

### 1. Subjects

DNA was extracted from peripheral blood of the Spanish ASD samples (360 trios; unaffected parents and affected proband) using the GentraPuregene blood kit (Qiagen Inc., Valencia, CA, USA). Subjects from Santiago (N = 136) were recruited from Complexo Hospitalario Universitario de Santiago de Compostela and Galician ASD organizations. Subjects from Madrid (N = 224) were recruited as part of AMITEA program at the Child and Adolescent Department of Psychiatry, Hospital General Universitario Gregorio Marañón. Only individuals 3 years old or older were included. All participants had a clinical diagnosis of ASC made by trained pediatric neurologists or psychiatrists based on the Diagnostic and Statistical Manual of Mental Disorders, Fourth Edition Text Revision and Fifth Edition (DSM-IV-TR and DSM-5) criteria. The Autism Diagnostic Observation Schedule (ADOS) and the Autism Diagnostic Interview-Revised (ADI-R) were also administered when necessary. Informed consent signed by each participating subject or legal guardian and approval from the corresponding Research Ethics Committee were obtained before the start of the study. All participants, parents or legal representatives provided written informed consent at enrollment and the study was conducted according to the declaration of Helsinki.

### 2. Sample quality control and DNMs detection

#### I. Data processing and annotation

WES of DNA extracted from the Spanish 360 trios was performed by the Autism Sequencing Consortium (ASC) (https://genome.emory.edu/ASC/)^21^. One multi-sample VCF with the raw results was retrieved from the ASC. Individual files containing coding variants per individual were obtained using bcftools and were annotated using SnpEff (Genomic variant annotations and functional effect prediction toolbox) version 4.3T (http://snpeff.sourceforge.net/)-.

#### II. Sample specific quality control

To check family relationships in the Spanish cohort (360 trios), information of Mendelian error counts was obtained using the “--mendel” option available in VCFtools (http://vcftools.sourceforge.net/). Samples whose Mendelian errors significantly deviated from the expectation were not considered for subsequent analysis.

To identify discrepancy between nominal designed and genetically determined sex the “--sexcheck” option in PLINK was used to infer correct sex from genotypes on chromosome X and Y.

Finally, to identify outlier samples in the Spanish cohort (360 trios), the “pseq i-stats” command in PLINK was used. Samples in which any of the following parameters: count of alternate, minor, heterozygous genotypes, number of called variants or genotyping rate deviated more than 4 SD from the mean were eliminated. Therefore, the whole trio was dropped if any member was considered an outlier.

The samples of the Spanish cohort (360 trios) that passed all the quality controls mentioned above were the same as those included in Satterstrom et al.^21^.

#### III. DNMs detection

To detect DNMs in the Spanish cohort (360 trios), defined as those mutations that are strictly present in probands and not in parents, the filtering options published by Lim et al. were employed^18^. In this study, variants classified as PZMs were resequenced by three different sequencing technologies reaching a high validation rate (87%-97%).

Briefly, we define DNMs as those variants whose genotypes were 1/0 or 1/1 in probands and 0/0 in parents. Then, variants with GQ ≥ 20 and alternate read depth ≥ 7 were considered. Variants that present two or more alleles in the ExAC database (http://exac.broadinstitute.org/) were filtered out. In addition, we filtered out variants that were less than 20 base pairs apart from each other to reduce false positives, and variants whose RVIS (Residual Variation Intolerance Score) retrieved from ExAC was higher than 75%. RVIS is designed to rank genes in terms of whether they have more or less common functional genetic variation relative to the genome-wide expectation given the amount of apparently neutral variation the gene has. Intolerant genes are more likely to be better candidates in NDDs. Thus, RVIS values represented as percentiles reflect the relative rank of the genes, with those genes above 75th percentile being the most intolerant.

SnpEff was employed to classify exonic variants according to the definition of their predictive impact: high, moderate and low impact on the canonical transcript. Low impact variants included silent mutations, moderate impact variants included missense mutations and high impact included splicing and nonsense mutations. Only base substitutions were considered so frameshift variants were filtered out.

Two different *in silico* prediction tools (CADD and SIFT) were used to classify missense mutations. Probably damaging mutations were those predicted as damaging by SIFT and variants with CADD score > 20 **(Additional file 1; Table S1)**.

Finally, DNMs were classified as germinal or PZMs based on the AAF (number of alternate reads / (total number of reference + alternate reads)). DNMs with an AAF ≥ 0.40 were classified as germinal and DNMs with an AAF < 0.40 were classified as PZMs^18^.

90 samples from the Spanish cohort (360 trios) were already analyzed by Lim et al.^18^ and they were used as positive controls to check if the detection of PZMs in the Spanish cohort was accurately made. Therefore, it was proved that most of DNMs were accurately detected and classified as germinal or PZMs **(Additional file 1; Tables S1, S2)**.

For the majority of the analysis, we used a dataset called “combined cohort” (N = 2171) which includes the non-synonymous DNMs detected in the 360 Spanish trios plus the non-synonymous DNMs identified in individuals with ASD sequenced by the ASC and published previously. Duplicated variants in both cohorts were eliminated (Supplementary table 3 of Lim et al.)^18^ **(Additional file 1; Table S1, S3)**. For some analysis, we also defined a control cohort of healthy siblings published by the ASC (same sequencing depth and variant calling procedures than the probands of the Spanish cohort) (N = 288)^181^ **(Additional file 1; Table S4)**.

### 3. Transmission and De *novo* Association Test (TADA-Denovo)

TADA-Denovo (http://www.compgen.pitt.edu/TADA/TADA_guide.html#tada-analysis-of-de-novo-data-tada-denovo) was run to discover and to prioritize ASD risk genes for both DNMs (germinal and PZMs) in the Spanish cohort (N = 360) **(Additional file 1; Table S1)** and in the combined dataset (N = 2171) **(Additional file 1; Table S3)**. TADA takes into account the mutational burden of the genes as well as the multiple mutational classes^22^. TADA was independently run in two different gene-sets for the Spanish cohort (genes harboring PZMs (PZMs genes) in the Spanish cohort = 105; genes harboring germinal DNMs (Germinal genes) in the Spanish cohort = 181) and for the combined cohort (PZMs genes in the combined cohort = 362; germinal genes in the combined cohort = 1210) **(Additional file 2; Tables S5, S6, S7 and S8)**. The control cohort (N = 288) was employed to set up and to estimate the parameters needed by TADA^18^ **(Additional file 1; Table S4)**.

Two classes of DNMs were included in the analysis: LoF and probably damaging missense mutations. To set up mutational rates for each mutational category, we used the per gene mutation rates table data computed by Samocha et al.^23^ and then, the following formula was applied to calibrate them: LoF; (nonsense+splice) x (syn_obs_/syn_exp_) and probably damaging missense; missense x (N_prob.damaging_ /N_allmissense_) x (syn_obs_/syn_exp_). Syn_obs_ is the observed number of synonymous mutations in the control cohort of unaffected siblings (N = 119)^18^, and syn_exp_ is the expected number of synonymous DNMs in the same cohort calculated from the sum of per-gene synonymous DNMs rates (2*n*μ) (N = 79.04). N_prob.damaging_ is the number of probably damaging missense mutations in the control cohort (N = 212) and N _allmissense_ were the total of missense mutations in the control cohort (N = 296). To estimate the relative risk (γ) for each mutational category, we calculated the burden (ƛ) of mutations of each type in cases (Spanish cohort) over controls (LoF = 2.21; probably damaging missense = 1.36). Then we applied the following formula to calculate relative risk: ɣ = 1+(ƛ-1)/ *π*, where *π*, the fraction of risk genes, was set as 0.05 (the default parameter). Finally, by running TADA-Denovo with the parameters described above, uncorrected p-values for each gene were calculated obtaining null distributions (N repetitions = 10,000). TADA-Denovo computes BF (Bayesian Factor) to each gene. To determine an appropriate threshold that allows declaring a “significant gene”, TADA uses the Bayesian FDR approach to control for the rate of false discoveries. q-values for each gene were calculated using the Bayesian FDR approach provided by TADA. Manhattan plots which show the results of TADA p-values (−log10) for the combined cohort (PZMs and germinal mutations) were done with R package qqman^24^.

Genes with FDR < 0.1 (germinal genes) and FDR < 0.3 (PZMs genes) were classified according to SFARI criteria (https://gene.sfari.org/database/gene-scoring/). Moreover, OMIM database (Online Mendelian Inheritance in Man) (https://www.omim.org/) was consulted to look for Mendelian diseases related to these genes.

### 4. Gene-set enrichment analysis of PZMs and germinal mutations

Gene-set enrichment analyses of those genes carrying missense and nonsense DNMs (germinal and PZMs) was done by DNENRICH^25^. DNENRICH estimates the enrichment of DNMs within pre-defined groups of genes accounting for gene size, tri-nucleotide context and functional effect of the mutations. DNMs included in this analysis were germinal and PZMs identified in the Spanish cohort (germinal DNMs = 236; PZMs = 164) **(Additional file 3; Tables S9 and S10)**. For the analysis of the combined cohort, a subset of PZMs was created in order to ensure that the analyzed PZMs likely contribute to the phenotype (PZMs = 676) **(Additional file 1; Table S11)**. For that purpose, individuals with germinal mutations in ASD risk genes (SFARI scores 1 and 2) were eliminated from the PZMs dataset. Thus, germinal DNMs and the subset of PZMs from the combined cohort were used in the analysis (germinal DNMs = 2270; PZMs = 676) **(Additional file 3; Table S12 and S13)**. The analysis was also run independently in unaffected siblings using data previously published by the ASC (germinal DNMs = 780; PZMs = 239) **(Additional file 1; Table S14)**.

The gene name alias and the gene size matrix provided by DNENRICH were used as input files used in this analysis together with the following gene-sets: 1) FMRP target genes identified by Darnell et al.^26^ and downloaded from Genebook (http://zzz.bwh.harvard.edu/genebook/) (N = 788); 2) Genes included in the GO:0006325 chromatin organization (N = 723) (http://www.geneontology.org/); 3) Synaptic genes (N = 903)^27^; 4) Human orthologs of genes essential in mice (N = 2472)^28^; 5) CHD8 target genes in human mid fetal brain (N = 2725)^29^; 6) List of SFARI genes (N = 990) (https://gene.sfari.org/autdb/HG_Home); 7) LoF intolerant genes (pLi > 0.9) (N = 3230)^30,29^; 8) RBFOX target genes (N = 587)^31^; 9) miR-137 target genes (N = 428)^32^; 10) CELF-4 target genes (N = 954)^33^; 11) Allele biased genes in differentiating neurons (N = 802)^34^; 12) Known ID genes (N = 1547)^35^; 13) Intergenic and Intronic Brain Expressed Enhancers (BEE) (N = 673)^36^; 14) Genomic intervals surrounding known telencephalon genes scanned for enhancers (N = 79)^37^; and 15) miR-138 target genes (N = 255)^38^. Empirical p-values were obtained from one million permutations for each gene-set.

### 5. Gene Ontology Enrichment Analysis

An exploratory GO enrichment analysis was carried out using the Enrichr tool. This analysis allows studying if genes harboring germinal DNMs or PZMs are involved in different biological processes. To this aim, the combined dataset of germinal missense and nonsense DNMs and the subset of PZMs were employed (germinal genes = 1972; PZMs genes = 624) **(Additional file 3; Table S15)**. In addition, the REViGO tool (http://revigo.irb.hr/) was employed to visualize GO terms in semantic similarity-based scatterplots using SimRel as a semantic similarity measure. Thus, the top 30 enriched GO terms in each group (PZM *vs* germinal) were visualized using a modification of the R script provided by the REViGO online tool.

Network visualization of the top 50 enriched terms in each group of genes was performed with Enrichment Map, a Cytoscape (v.3.6.1) plugin for functional enrichment visualization^39^. Each node represents a gene-set (GO term) and the size of the node is proportional to the number of genes participating in the GO term (overlap coefficient). Nodes were considered as connected when the overlap coefficient was greater than 0.7 and edge-width represents the overlap between gene-sets. The border-width of each node represents the corresponding p-value for each GO term.

### 6. Expression cell-type enrichment analysis and expression analysis across brain regions and developmental periods

Expression Weighted Cell-type Enrichment (EWCE) method (https://github.com/NathanSkene/EWCE) was used to explore whether genes harboring germinal DNMs (N = 1972) and genes harboring PZMs (N = 624) **(Additional file 3; Table S15)** were differentially expressed across several neuronal cell types. EWCE involves testing whether the given genes in a target list have higher levels of expression in a given cell type compared to what is expected by chance. Brain single-cell transcriptomic data from Karolinska Institute (ctd_allKI) was used for the EWCE analysis. Brain regions included in the KI mouse super dataset are the neocortex, hippocampus, hypothalamus, striatum, and midbrain, as well as samples enriched for oligodendrocytes, dopaminergic neurons and cortical parvalbumin interneurons (total cells = 9970). Background gene-set comprises all human-mice orthologous. Probability distribution for our gene lists was calculated by randomly sampling 100000 genes from the background set controlling for transcript length and GC content. Bootstrapping function was then applied on level 1 annotation.

pSI (specificity index statistic), an R package, was employed to study the expression of genes harboring PZM and germinal mutations across different brain regions and neurodevelopmental periods^40,41^. Lists of specifically expressed human genes (human.rda) obtained from BrainSpan data (gene-sets for 6 brain regions and gene-sets for 10 developmental periods) were employed. The Fisher iteration test included in the pSI package was used to analyze if the listed genes harboring PZM and germinal mutations were significantly overrepresented. Brain areas significantly enriched with PZMs or germinal genes in specific developmental periods (p-adjusted value < 0.05) were represented as a matrix.

## RESULTS

### 1. Transmission and De *novo* Association Test (TADA-Denovo)

Transmission and De *novo* Association test (TADA) assesses if a gene is affecting ASD risk based on several parameters: the gene mutation rate, the recurrence of DNMs in the gene and the severity of the mutations^22^. Thus, TADA-Denovo analysis was independently run in both datasets (germinal and PZMs genes). The main aim of TADA-Denovo is to identify those genes that could be differentially involved in ASD etiology depending on the type of DNMs harbored by them (germinal or PZMs). We focused the analysis on damaging mutations (LoF and likely pathogenic missense variants) to increase the likelihood of finding “strong” candidate genes. First, the set of genes from the Spanish cohort (360 trios) (germinal genes = 181; PZMs genes = 105) was analyzed. The analysis of the germinal gene list identified 12 genes with an FDR < 0.3 **(Table 1 and Additional file 2; Table S16)** including 3 genes (*SCN2A, ARID1B* and *CHD8*) with an FDR < 0.1. The analysis of the PZMs gene list identified 13 genes with an FDR < 0.3 **(Table 2 and Additional file 2; Table S17)** of which 4 genes (*KMT2C, FRG1, GRIN2B* and *MAP2K3*) had an FDR < 0.1.

**Table 1.** ASD risk genes carrying germinal mutations in the Spanish cohort. p-values and q-values were obtained after running TADA-Denovo using germinal DNMs from the Spanish ASD cohort (N=360). Only genes with q-values <0.3 are shown.

| Genes | q-value | p-value |
| --- | --- | --- |
| **SCN2A** | 0.004 | $2.76 \times 10^{-7}$ |
| **ARID1B** | 0.050 | $2.76 \times 10^{-6}$ |
| **CHD8** | 0.066 | $3.31 \times 10^{-6}$ |
| FIG4 | 0.103 | $9.39 \times 10^{-5}$ |
| RBM15 | 0.126 | $1.16 \times 10^{-5}$ |
| HUWE1 | 0.148 | $3.54 \times 10^{-5}$ |
| KIAA1107 | 0.188 | $5.08 \times 10^{-5}$ |
| VWAS5B1 | 0.218 | $5.08 \times 10^{-5}$ |
| EMCN | 0.242 | $6.13 \times 10^{-5}$ |
| SH2B2 | 0.262 | $7.90 \times 10^{-5}$ |
| ASMT | 0.277 | $9.34 \times 10^{-5}$ |
| MYLK4 | 0.291 | $9.67 \times 10^{-5}$ |

**Table 2.** ASD risk genes carrying PZMs in the Spanish cohort. p-values and q-values were obtained after running TADA-Denovo using PZMs from the Spanish ASD cohort (N=360). Only genes with q-values <0.3 are shown.

| Genes | q-value | p-value |
| --- | --- | --- |
| KMT2C | 0.001 | $4.76 \times 10^{-7}$ |
| FRG1 | 0.015 | $4.46 \times 10^{-7}$ |
| GRIN2B | 0.040 | $4.76 \times 10^{-7}$ |
| MAP2K3 | 0.08 | $6.67 \times 10^{-6}$ |
| SRGAP2 | 0.106 | $7.62 \times 10^{-6}$ |
| MBD6 | 0.124 | $7.62 \times 10^{-6}$ |
| POTEB2 | 0.168 | $2.57 \times 10^{-5}$ |
| CALML6 | 0.200 | $2.67 \times 10^{-5}$ |
| PRDX6 | 0.226 | $3.05 \times 10^{-5}$ |
| SSR2 | 0.245 | $3.14 \times 10^{-5}$ |
| VEGFA | 0.264 | $4 \times 10^{-5}$ |
| CANX | 0.278 | 0.0001 |
| ZNF276 | 0.290 | 0.00012 |

In the combined cohort (genes from the Spanish cohort plus genes from the Lim et al. publication ^18^(germinal genes = 1210; PZMs genes = 362) TADA identified 34 genes with an FDR < 0.1 **(Table 3 and Figure 1a)** and 103 genes with an FDR < 0.3 **(Additional file 2; Table S18)**. Three of the genes (*SCN2A, ARID1B, CHD8*) with germinal DNMs were prioritized (FDR < 0.1) both in the combined cohort and in the Spanish cohort when TADA. Analysis of PZMs genes in the combined cohort identified three genes (*FRG1, KMT2C* and *NFIA*) with an FDR < 0.1, and 14 genes with an FDR < 0.3 **(Table 4 and Figure 1b; Additional file 2; Table S19)**. Only two genes, *KMT2C* and *FRG1*, have remained significant after FDR correction (< 0.1) in both the Spanish and the combined cohort.

**Table 3.** ASD risk genes carrying germinal mutations in the combined cohort. p-values and q-values were obtained after running TADA-Denovo in the combined cohort (N=2103). Genes with q-values <0.1 are shown.

| Gene | q-value | p-value |
| --- | --- | --- |
| **SCN2A** | $5.04 \times 10^{-12}$ | $4.13 \times 10^{-8}$ |
| **CHD8** | $2.40 \times 10^{-5}$ | $4.13 \times 10^{-8}$ |
| **ARID1B** | $5.36 \times 10^{-5}$ | $4.13 \times 10^{-8}$ |
| SLC6A1 | 0.00015 | $2.48 \times 10^{-7}$ |
| SYNGAP1 | 0.0005 | $6.61 \times 10^{-7}$ |
| KDM5B | 0.0008 | $8.26 \times 10^{-7}$ |
| SUV420H1 | 0.002 | $5.37 \times 10^{-6}$ |
| TRIP12 | 0.003 | $5.79 \times 10^{-6}$ |
| PTEN | 0.004 | $1.14 \times 10^{-5}$ |
| KATNAL2 | 0.008 | $5.01 \times 10^{-5}$ |
| NRXN1 | 0.012 | $5.79 \times 10^{-5}$ |
| CREBBP | 0.02 | $5.92 \times 10^{-5}$ |
| CELF4 | 0.02 | $6.09 \times 10^{-5}$ |
| STXBP1 | 0.02 | $6.48 \times 10^{-5}$ |
| DYRK1A | 0.02 | $7.09 \times 10^{-5}$ |
| CHD2 | 0.03 | 0.0001 |
| ANK2 | 0.03 | 0.0001 |
| WDFY3 | 0.03 | 0.0001 |
| UNC80 | 0.04 | 0.0002 |
| CLASP1 | 0.04 | 0.0002 |
| TMEM39B | 0.05 | 0.0002 |
| PRKAR1B | 0.05 | 0.0002 |
| USP45 | 0.05 | 0.0003 |
| NUAK1 | 0.06 | 0.0004 |
| NAA15 | 0.06 | 0.0004 |
| FOXP1 | 0.07 | 0.0004 |
| ZC3H11A | 0.07 | 0.0004 |
| DPP3 | 0.07 | 0.0005 |
| PRKDC | 0.08 | 0.0005 |
| ATP1A1 | 0.08 | 0.0005 |
| LRP5 | 0.09 | 0.0005 |
| SLC12A3 | 0.09 | 0.0006 |
| FBXO18 | 0.096 | 0.0006 |
| PTK7 | 0.0999 | 0.0007 |

**Figure 1.**
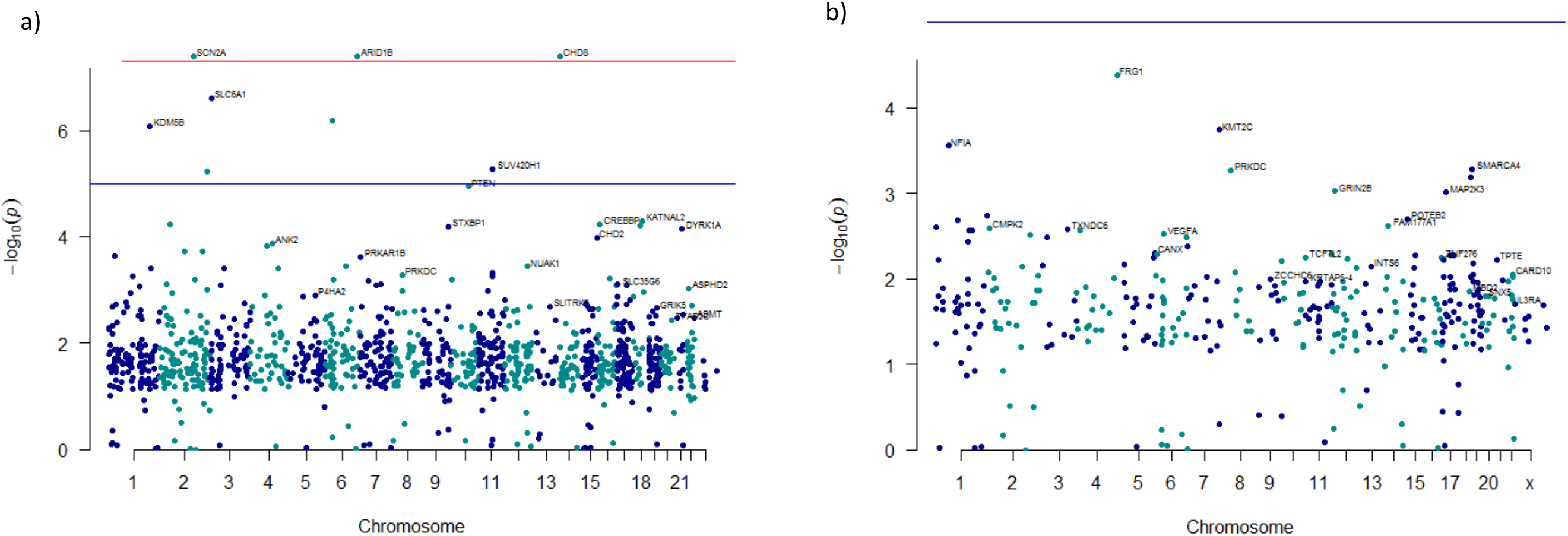
Manhattan plots depicting the ASD risk genes prioritized by TADA-Denovo. (chromosome and log10 p-value for each gene are represented in axis x and y). a) p-values were obtained from analysis of germinal mutations in the combined cohort using TADA-Denovo. Red line represents the p-value < 1 x 10^−8^ and blue line p-value < 1 x 10 ^−5^. b) p-values were obtained from analysis of PZMs in the combined cohort using TADA-Denovo. Blue line represents p-value < 1 x 10 ^−5^.

**Table 4.** ASD risk genes carrying PZMs in the combined cohort. p-values and q-values were obtained after running TADA-Denovo in the combined cohort (N=2103). Genes with q-values < 0.3 are shown.

| Gene | q-value | p-value |
| --- | --- | --- |
| *FRG1* | 0.04 | 4.14 x 10<sup>-3</sup> |
| *KMT2C* | 0.07 | 0.00018 |
| *NFIA* | 0.09 | 0.00028 |
| *SMARCA4* | 0.12 | 0.00052 |
| *PRKDC* | 0.13 | 0.00055 |
| *KLF16* | 0.15 | 0.00064 |
| *GRIN2B* | 0.17 | 0.00095 |
| *MAP2K3* | 0.18 | 0.00098 |
| *HNRNPU* | 0.21 | 0.0019 |
| *POTEB2* | 0.23 | 0.002 |
| *RNPC3* | 0.25 | 0.002 |
| *FAM177A1* | 0.27 | 0.002 |
| *CALML6* | 0.28 | 0.002 |
| *CMPK2* | 0.3 | 0.003 |

A total of 17 genes (50%) from the set of germinal genes in the combined cohort (34 genes, FDR < 0.1) were identified as “high confidence” or “strong” ASD candidates following SFARI Gene scoring criteria (scores 1, 2, 1s and 2s). In addition, 11 of these genes (64,70%) have shown an FDR < 0.1 in previous TADA analysis^6^. Moreover, 10 of the remaining genes identified by TADA (FDR < 0.1) are included in SFARI gene lists (scores 3, 4 and 5) and 5 of the genes identified by TADA were reported in relation with another disease (not ASD) by OMIM database **(Additional file 4; Table S20)**.

PZM analysis has shown association of *KMT2C* (SFARI score s2) as well as other 3 genes FDR < 0.1). It is worth to note that *NFIA* has been previously reported as a plausible candidate gene in ASD (SFARI score 4) but this is the first time that *FRG1* is reported in ASD. *SMARCA4, PRKDC, KLF16, GRIN2B* and *HNRNPU* (SFARI score 3 and 4) were among those plausible ASD candidate genes previously identified with an FDR value between 0.1 and 0.3. GRIN2B was previously reported by SFARI as a strong ASD risk gene (score 1) **(Additional file 4; Table S21)**.

### 2. Gene-set enrichment analysis of PZMs and germinal mutations

DNENRICH was run to estimate a statistical significance of enrichment for germinal and PZMs within previously ASD and NDDs associated gene-sets. Synonymous mutations were excluded from the analysis because they are unlikely to contribute to ASD phenotype and only nonsense and missense mutations were considered. First, gene-set enrichment analysis was performed using the list of genes and DNMs (germinal and PZMs) from the Spanish cohort **(Additional file 3; Tables S9 and S10)** against several background gene lists **(see Methods)**. Our results indicate that germinal genes shown enrichment in several gene-sets (germinal genes = 228, germinal DNMs = 236): FMRP target genes (p-value = 0.003), known ID genes (p-value = 0.0073), LoF intolerant genes (p-value = 0.002), SFARI genes (p-value = 1 x 10^−7^) and genes involved in chromatin organization (p-value = 0.00018) **(Table 5)**. However, only the LoF intolerant gene-set has shown association with the list of PZMs genes (PZMs genes = 155, PZMs = 164) **(Table 6)**.

**Table 5.** Results of the gene-set enrichment analysis for the list of genes harboring germinal mutations (Spanish cohort).

| Gene-sets | p-value | Observed mutations | Expected mutations |
| --- | --- | --- | --- |
| Allele biased genes in differentiating neurons | 0.6 | 12 | 12.502 |
| CELF4 targets | 1 | 0 | 0.05078 |
| Essential genes | 0.0709 | 49 | 39.968 |
| FMRP targets | 0.00013 | 39 | 20.9788 |
| Genomic intervals surrounding known telencephalon genes | 0.5386 | 1 | 0.771873 |
| CHD8 targets | 0.226 | 39 | 34.4983 |
| Known ID genes | 0.0073 | 40 | 27.0286 |
| LoF intolerant genes | 0.0018 | 79 | 58.9156 |
| SFARI genes | $1 \times 10^{-6}$ | 49 | 20.9601 |
| Synaptic genes | 0.3592 | 14 | 12.4002 |
| Genes involved in chromatin organization | 0.00018 | 24 | 10.7377 |
| mir137 targets | 0.3251 | 8 | 6.49354 |
| mir128 targets | 1 | 0 | 0.022676 |
| RBFOX targets | 1 | 0 | 0.055409 |

**Table 6.** Results of the gene-set enrichment analysis for the list of genes harboring PZM (Spanish cohort).

| Gene-sets | p-value | Observed mutations | Expected mutations |
| --- | --- | --- | --- |
| Allele biased genes in differentiating neurons | 0.9727 | 4 | 8.48746 |
| CELF4 targets | 1 | 0 | 0.034487 |
| Essential genes | 0.2340 | 31 | 27.1352 |
| FMRP targets | 0.0764 | 20 | 14.2422 |
| Genomic intervals surrounding known telencephalon genes scanned for enhancers | 0.0977 | 2 | 0.524631 |
| CHD8 targets | 0.0884 | 30 | 23.4115 |
| Known ID genes | 0.7554 | 16 | 18.3456 |
| LoF intolerant genes | 0.0283 | 51 | 39.9973 |
| Sfari genes | 0.0764 | 20 | 14.2361 |
| Synaptic genes | 0.3339 | 10 | 8.41702 |
| Genes involved in chromatin organization | 0.0625 | 12 | 7.28793 |
| mir137 targets | 0.8198 | 3 | 4.40692 |
| mir128 targets | 1 | 0 | 0.015316 |
| RBFOX targets | 1 | 0 | 0.037786 |

DNENRICH analysis in the combined cohort demonstrated enrichment for several gene-sets for both germinal and PZMs genes: chromatin organization, SFARI genes, LoF intolerant genes, CHD8 target genes and essential genes. In addition, the germinal gene list (germinal genes = 1972, germinal DNMs = 2270) showed enrichment for FMRP target genes (p-value = 1x 10^−6^), known ID genes (p-value = 1 x 10^−6^) and synaptic genes (p-value = 4x 10^−6^) **(Table 7, Figure 2)**. PZMs genes (PZMs genes = 624, PZMs = 676), have only shown association in the case of the miR-137 target gene-set (p-value = 0.0019) **(Table 8, Figure 2)**. The same analysis was performed using the list of germinal genes from unaffected siblings (germinal genes= 744, germinal DNMs = 780; PZMs genes= 237, PZMs = 239) **(Additional file 1; Table S14)**. A significant enrichment was identified only with the FMRP targets gene-set (data not shown).

**Table 7.**
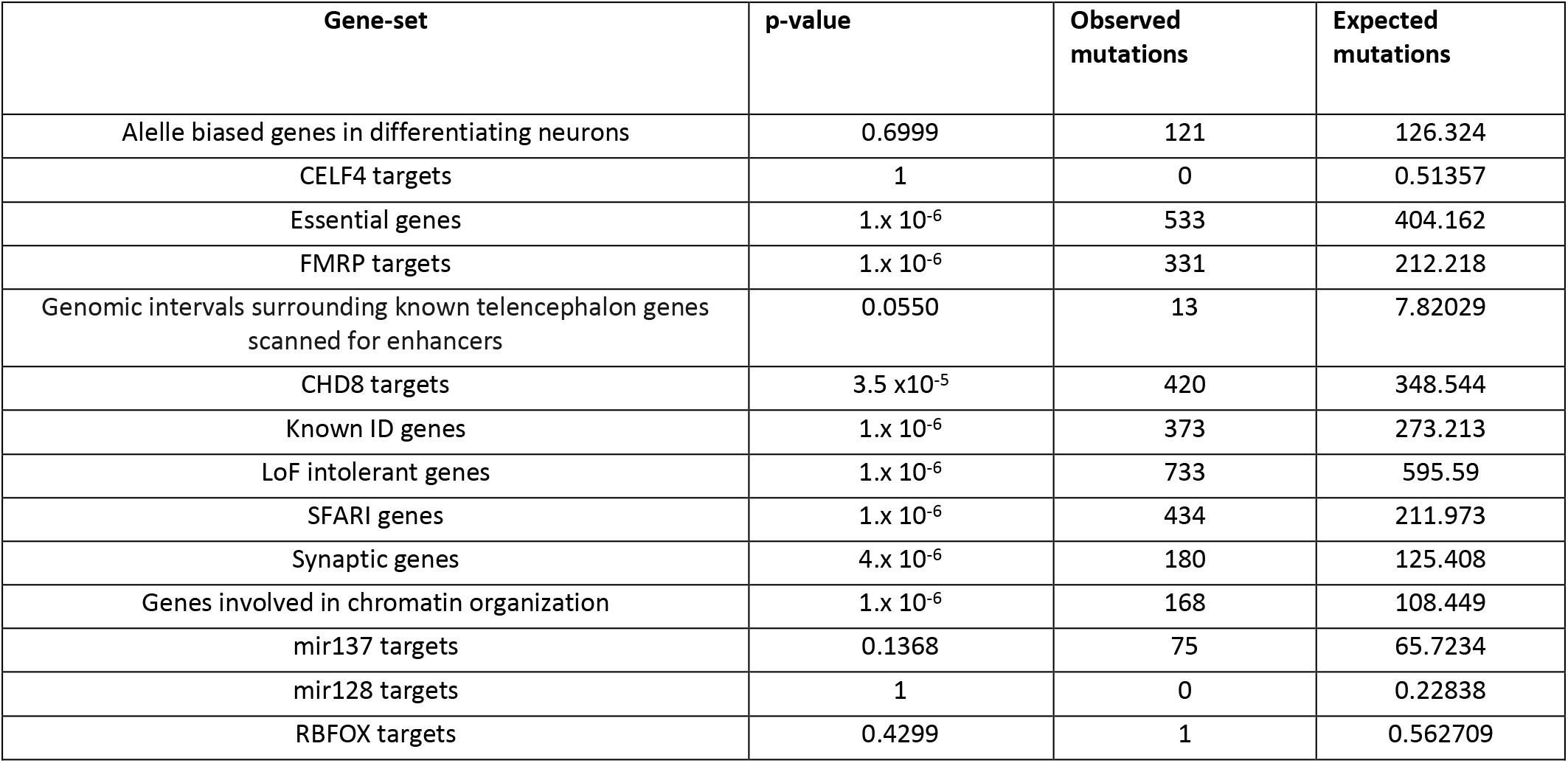
Results of the gene-set enrichment analysis for the list of genes harboring germinal mutations (combined cohort).

| Gene-set | p-value | Observed mutations | Expected mutations |
| --- | --- | --- | --- |
| Allele biased genes in differentiating neurons | 0.6999 | 121 | 126.324 |
| CELF4 targets | 1 | 0 | 0.51357 |
| Essential genes | $1. \times 10^{-6}$ | 533 | 404.162 |
| FMRP targets | $1. \times 10^{-6}$ | 331 | 212.218 |
| Genomic intervals surrounding known telencephalon genes scanned for enhancers | 0.0550 | 13 | 7.82029 |
| CHD8 targets | $3.5 \times 10^{-5}$ | 420 | 348.544 |
| Known ID genes | $1. \times 10^{-6}$ | 373 | 273.213 |
| LoF intolerant genes | $1. \times 10^{-6}$ | 733 | 595.59 |
| SFARI genes | $1. \times 10^{-6}$ | 434 | 211.973 |
| Synaptic genes | $4. \times 10^{-6}$ | 180 | 125.408 |
| Genes involved in chromatin organization | $1. \times 10^{-6}$ | 168 | 108.449 |
| mir137 targets | 0.1368 | 75 | 65.7234 |
| mir128 targets | 1 | 0 | 0.22838 |
| RBFOX targets | 0.4299 | 1 | 0.562709 |

**Figure 2.**
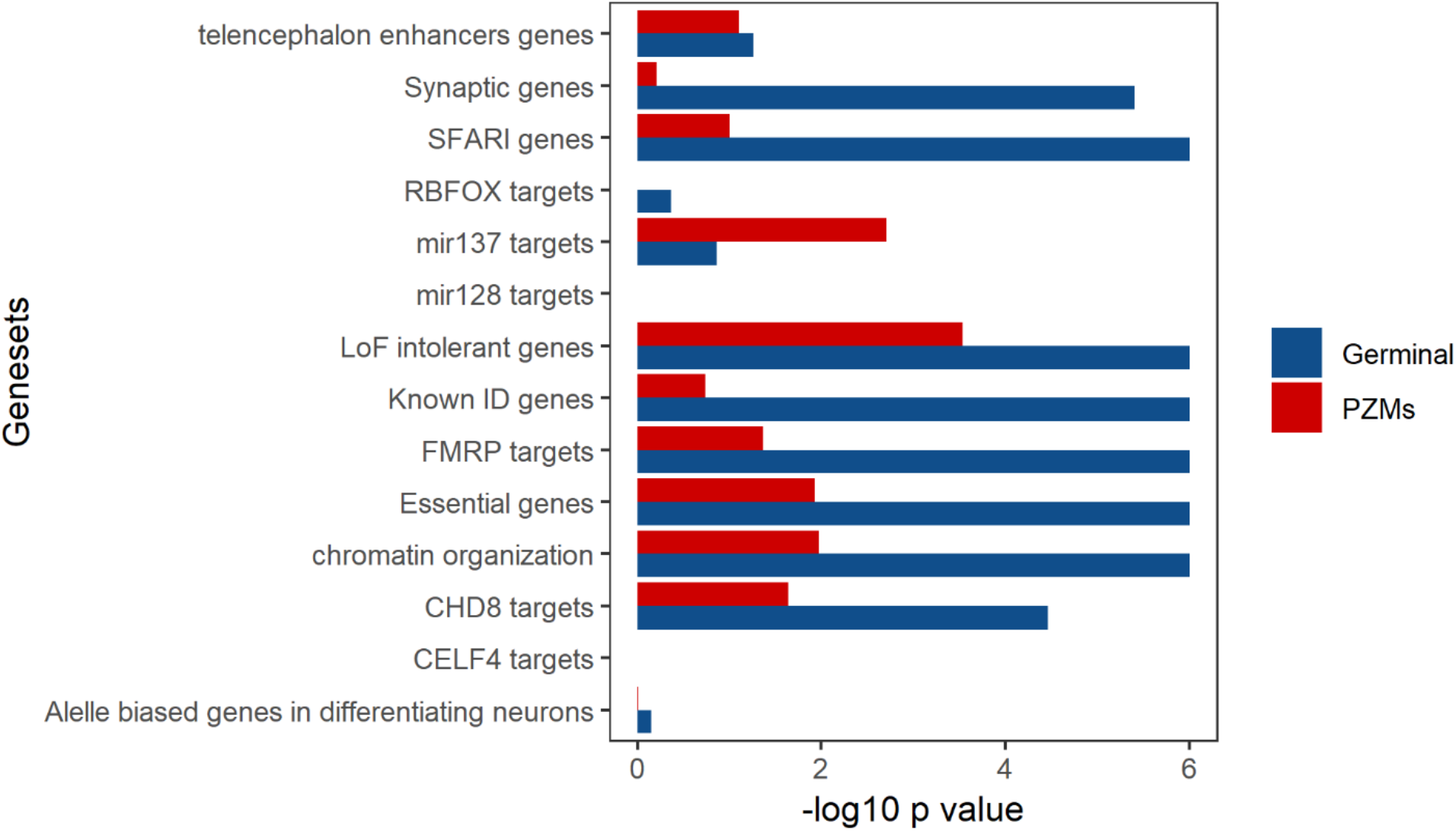
Gene-set enrichment analysis using germinal and PZM from the combined cohort. Gene-set enrichment analysis was done with DNENRICH. −log10 p-value for each gene-set is shown for each type of mutation and tested gene-set.

**Table 8.** Results of the gene-set enrichment analysis for the list of genes harboring PZM (combined cohort).

| Gene-set | p-value | Observed mutations | Expected mutations |
| --- | --- | --- | --- |
| Allele biased genes in differentiating neurons | 0.9744 | 26 | 36.4938 |
| CELF4 targets | 1 | 0 | 0.147914 |
| Essential genes | 0.0118 | 140 | 116.895 |
| FMRP targets | 0.0425 | 75 | 61.4294 |
| Genomic intervals surrounding known telencephalon genes scanned for enhancers | 0.0796 | 5 | 2.2672 |
| CHD8 targets | 0.0228 | 120 | 100.751 |
| Known ID genes | 0.1840 | 87 | 79.0127 |
| LoF intolerant genes | 0.0003 | 212 | 172.214 |
| SFARI genes | $1 \times 10^{-7}$ | 127 | 61.2987 |
| Synaptic genes | 0.6109 | 35 | 36.2813 |
| Genes involved in chromatin organization | 0.0106 | 45 | 31.3424 |
| mir137 targets | 0.0019 | 33 | 19.0234 |
| mir128 targets | 1 | 0 | 0.066195 |
| RBFOX targets | 1 | 0 | 0.162842 |

### 3. Gene Ontology Enrichment Analysis

GO enrichment analysis revealed remarkable differences between germinal and PZMs gene lists from the combined cohort **(Additional file 3; Table S15)**. The germinal set showed a significant enrichment in different GO terms related to synaptic function and transcription regulation. In particular, it is worth to note the association of GO terms related to ion transport: GO:0006814, Benjamini-Hochberg-corrected p[Pbh] = 0.005; GO0035725, Pbh = 0.005 and GO0006816, Pbh = 0.006 **(Additional file 4; Table S22, Figure 3a)**. The GO terms enriched in the subset of PZMs are related to regulation of gene expression, biosynthesis, differentiation or migration: GO0010629, p[Pbh] = 0.074; GO:2000113, p[Pbh] = 0.092; GO:0045652, p[Pbh] = 0.092; GO:0030336, p[Pbh] = 0.0995. **(Additional file 4; Table S23, Figure 3b)**.

**Figure 3.**
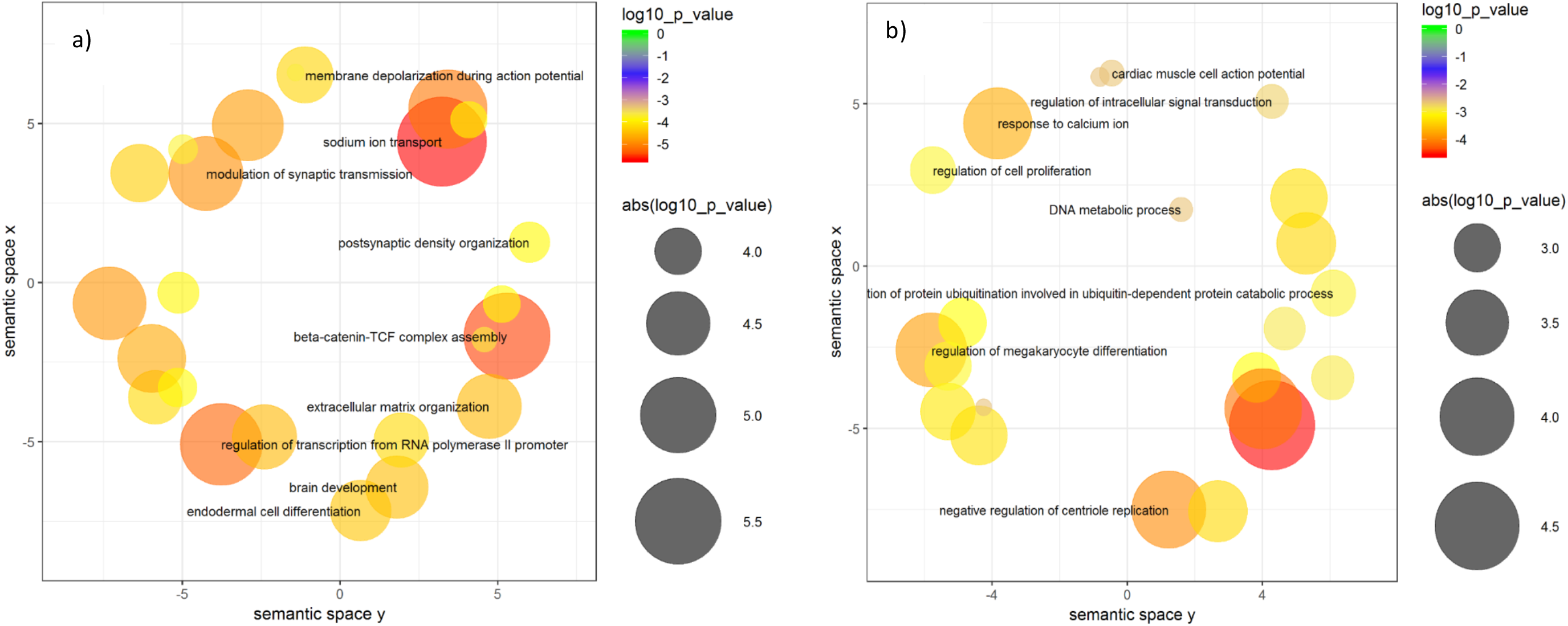
Scatterplots representing the top 30 significant biological processes (combined cohort). a) Top 30 biological processes enriched in genes harboring germinal DNM are shown. b) Top 30 biological processes enriched in genes harboring PZMs DNMs are shown.

GO enrichment analysis was depicted by semantically clustering the top 50 enriched terms for germinal and PZMs gene lists. The germinal gene list resulted in three differentiated clusters: neuron development and differentiation, synaptic functions, and chromatin modifications. The existence of a fourth cluster, which includes terms related to embryonic development, was also highlighted **(Figure 4a)**. In the case of PZMs genes, all the clusters were partially related to each other. However, we identified another cluster that includes terms related to the regulation of core processes (e.g, protein phosphorylation, regulation of growth, negative regulation of cellular biosynthetic processes, positive regulation of transcription DNA template). It is also important to highlight the GO terms related to neuron and embryonic development **(Figure 4b)**.

**Figure 4.**
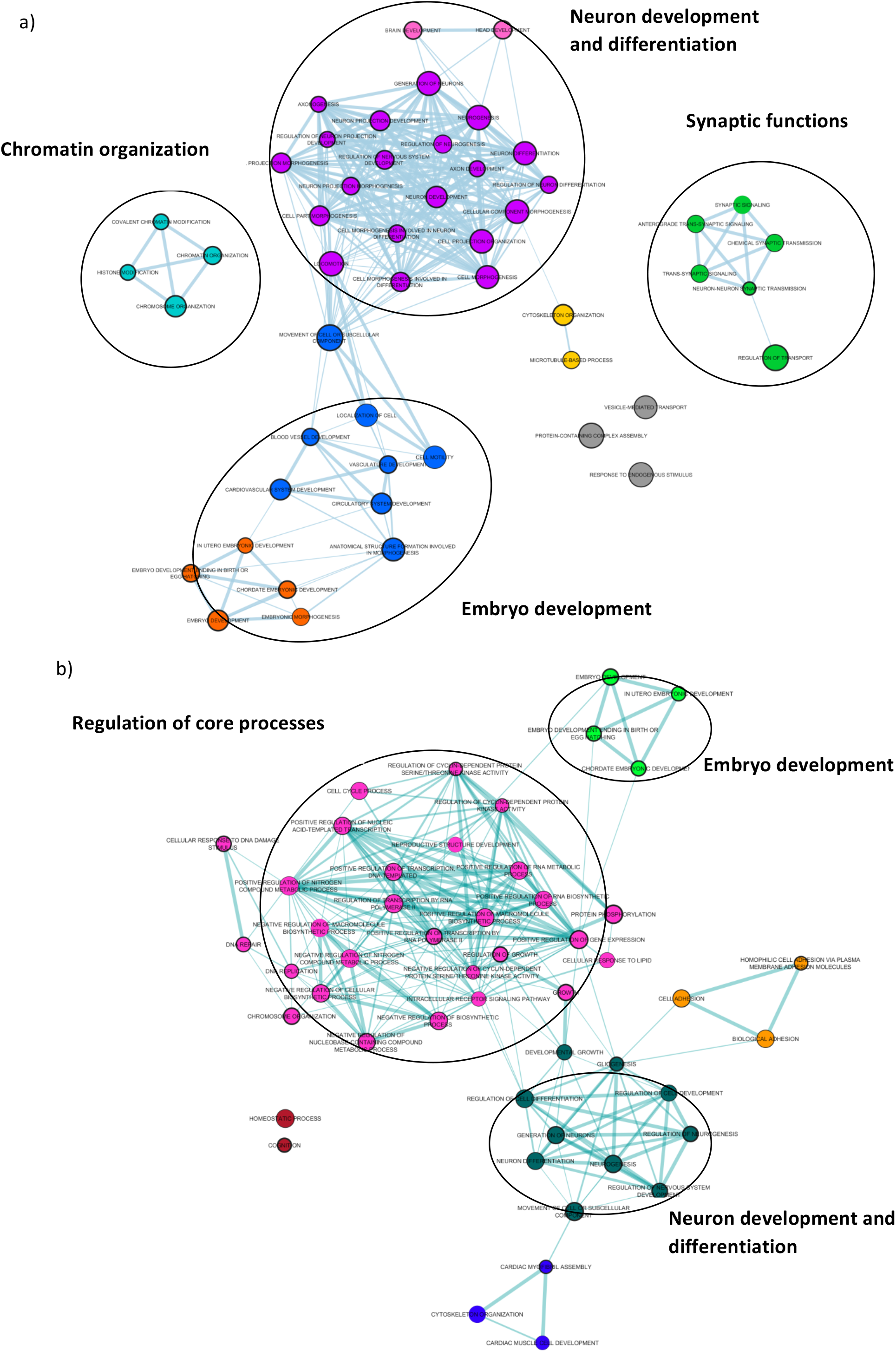
Visualization of the top 50 GO terms grouped by clusters of biological functions (DNMs obtained from the combined cohort) a) GO terms clusters for genes harboring germinal mutations. b) GO terms clusters for genes harboring PZMs.

### 4. Expression cell-type enrichment analysis and expression analysis across brain regions and developmental periods

First, we examined whether germinal genes or PZMs genes from the combined cohort, were differentially expressed in the transcriptome dataset corresponding to level 1 cell types. As expected, germinal genes were significantly enriched in several cell types **(Table 9; Figure 5)**. The most enriched cell types were those related to neurotransmission (dopaminergic neuroblast; p-value < 0.0001; embryonic dopaminergic neurons, p-value < 0.0001; embryonic GABAergic neurons, p-value < 0.0001; serotonergic neurons, p-value < 0.0001). The PZMs gene list showed enrichment for three different cell types: pyramidal CA1 neurons; p-value = 0.0066, pyramidal somatosensory (SS); p-value = 0.016, embryonic midbrain nucleus neurons; p-value = 0.02 **(Table 10, Figure 5)**.

**Table 9.** Top 10 cell enrichment in genes disrupted by germinal mutations in the combined cohort.

| Cell type | p | fold_change | sd_from_mean |
| --- | --- | --- | --- |
| Dopaminergic Neuroblast | <0.0001 | 1.17053835 | 4.27167674 |
| Embryonic Dopaminergic Neuron | <0.0001 | 1.175277135 | 5.113211893 |
| Embryonic GABAergic Neuron | <0.0001 | 1.11078582 | 4.626262187 |
| Neuroblasts | <0.0001 | 1.187690135 | 5.49208347 |
| pyramidal CA1 | <0.0001 | 1.132632722 | 5.365027275 |
| pyramidal SS | <0.0001 | 1.12556771 | 5.157499425 |
| Serotonergic Neuron | <0.0001 | 1.170544581 | 5.491769105 |
| Neural Progenitors | 0.0001 | 1.15212965 |  |
| Radial glia like cells | 0.0003 | 1.124574836 | 3.589901456 |
| Embryonic midbrain nucleus neurons | 0.0004 | 1.104053784 | 3.513053618 |

**Figure 5.**
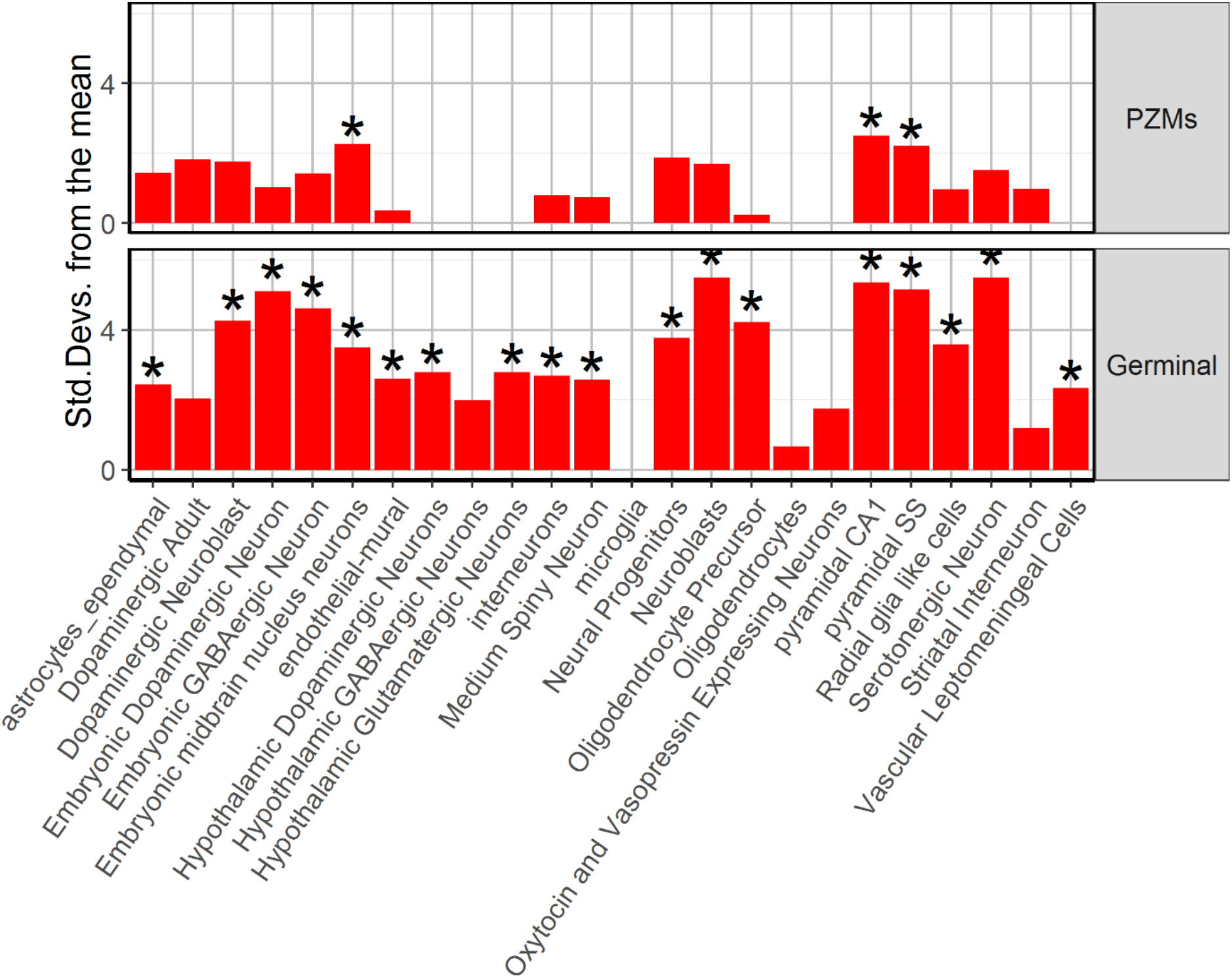
Expression Weighted Cell type Enrichment for PZM and germinal genes.

**Table 10.** Top 10 cell enrichment in genes disrupted by PZMs mutations in the combined cohort.

| Cell type | p | fold_change | sd_from_mean |
| --- | --- | --- | --- |
| pyramidal CA1 | 0.0066 | 1.116362092 | 2.509881972 |
| pyramidal SS | 0.0152 | 1.09832511 | 2.209046983 |
| Embryonic midbrain nucleus neurons | 0.0185 | 1.125338403 | 2.263702976 |
| Neural Progenitors | 0.0336 | 1.141154419 | 1.880862217 |
| Dopaminergic Adult | 0.0404 | 1.096578007 | 1.823257602 |
| Dopaminergic Neuroblast | 0.0474 | 1.130737104 | 1.762033343 |
| Neuroblasts | 0.0572 | 1.108135618 | 1.686197427 |
| Serotonergic Neuron | 0.067 | 1.090682473 | 1.525134485 |
| astrocytes_ependymal | 0.0802 | 1.123951233 | 1.432463223 |
| Embryonic GABAergic Neuron | 0.0827 | 1.062881294 | 1.417776316 |

To gain insight into the spatiotemporal distribution, we analyzed the expression of germinal and PZMs genes **(Additional file 3; Table S15)** across several brain regions and different neurodevelopmental periods obtained from BrainSpan. Germinal genes were significantly expressed in the cortex, striatum, cerebellum and amygdala in prenatal stages (early, early mid and late) **(Table 11, Figure 6a and 6c)**. PZMs genes were significantly expressed in the cortex during the early mid-fetal period. Although we did not find found a significant enrichment in other brain areas or neurodevelopmental periods for PZMs, p-values close to the significance threshold were found in cortex (early, late mid-fetal) and amygdala (late mid-fetal) **(Table 12, Figure 6b and 6d)**.

**Table 11.** Expression analysis of germinal targeted genes from the combined cohort in brain regions and developmental periods

| Brain region | Developmental period | p-value |
| --- | --- | --- |
| Cortex | Early Fetal | 0.003 |
| Cortex | Early Mid Fetal | 0.003 |
| Cortex | Young Adulthood | 0.003 |
| Striatum | Early Fetal | 0.008 |
| Cerebellum | Late Mid Fetal | 0.027 |
| Amygdala | Early Mid Fetal | 0.046 |

**Figure 6.**
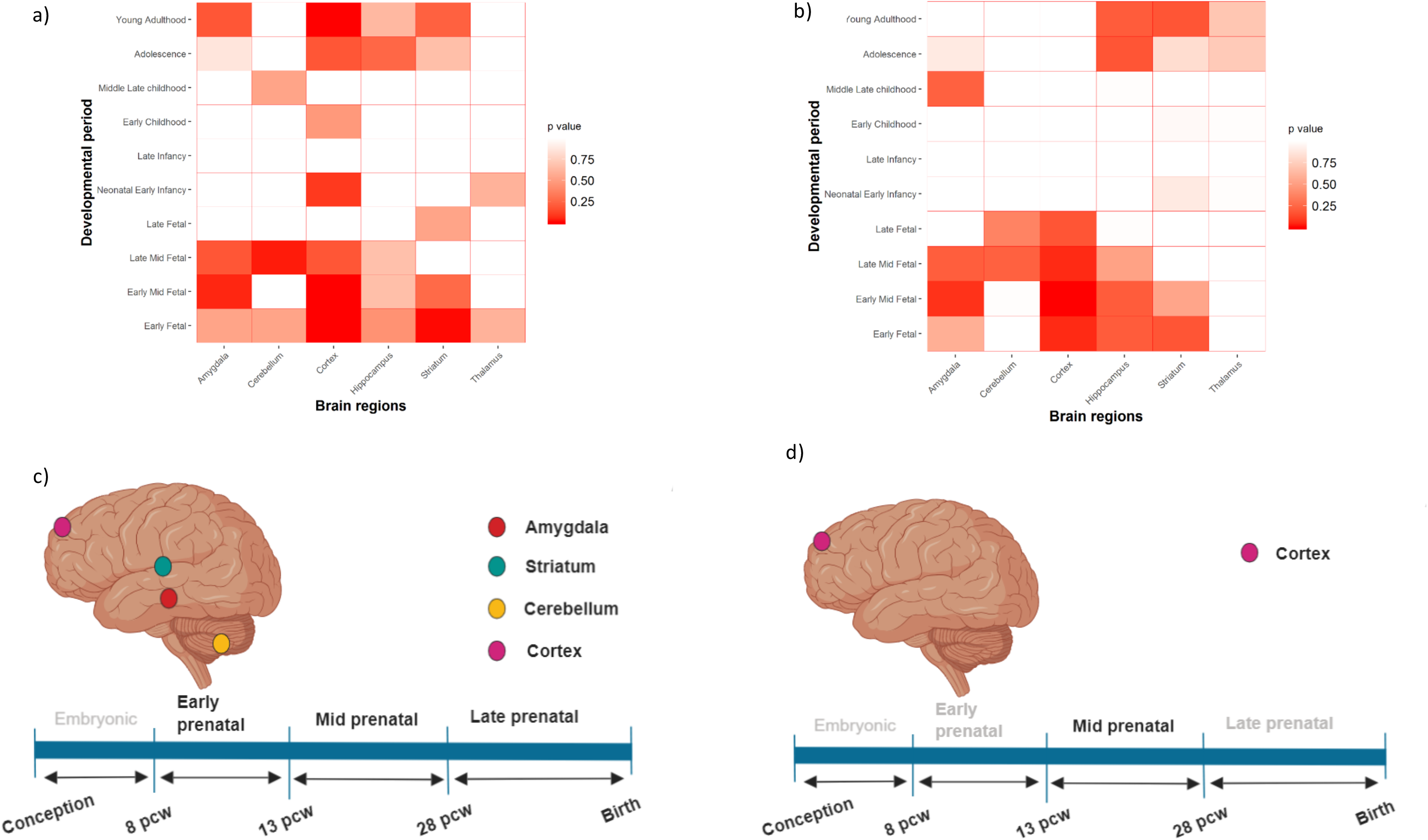
Expression analysis across developmental stages and brain areas for germinal and PZMs genes from the combined cohort. Expression gene sets were obtained from BrainSpan. a and c) Expression of germinal genes across different brain regions and developmental periods. b and d) Expression of PZMs genes across different brain regions and developmental periods

**Table 12.**
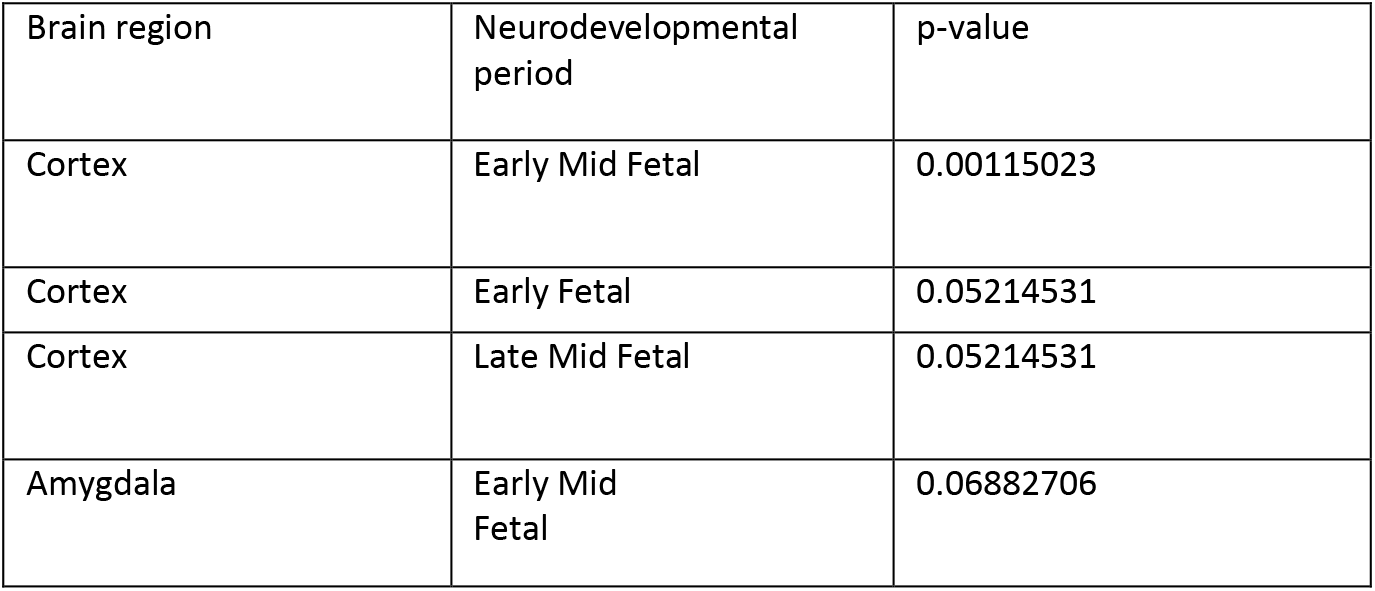
Expression analysis of PZMs targeted genes from the combined cohort in brain regions and developmental periods

| Brain region | Neurodevelopmental period | p-value |
| --- | --- | --- |
| Cortex | Early Mid Fetal | 0.00115023 |
| Cortex | Early Fetal | 0.05214531 |
| Cortex | Late Mid Fetal | 0.05214531 |
| Amygdala | Early Mid Fetal | 0.06882706 |

## DISCUSSION

ASD is a complex NDD, characterized by its clinical and genetic heterogeneity. Over 1000 genes are estimated to be involved in ASD pathogenesis but only a small number are functionally well characterized. Thus, it is widely believed that the vast majority of ASD genetic factors remain largely unknown^6^. In addition, DNMs are a strong source of genetic causality in ASD but they have been usually identified in a germinal state. Recent studies have suggested that some ASD candidate genes could carry more mosaic mutations than others^15–18^. However, the phenotypic effect of PZMs and its impact in clinical diagnosis is not completely understood. A recent study showed that brain malformations can occur when around 10% of peripheral blood cells harbor the mosaic mutation^42^. In addition, mosaic mutations could act as phenotype modifiers, resulting in patients who have less severe symptoms^43^.

Three genes were associated in the PZM analysis done by TADA (q-value < 0.1; *FRG1*, *KMT2C* and *NFIA*). *FRG1* has not been previously associated with NDDs while both, *KMT2C* and *NFIA*, have been previously reported as possible implicated in ASD/ID^44,45^. However, none of these genes was found to be associated to ASD when the analysis was performed in the list of germinal genes. These results point to the fact that different ASD risk genes could tend to differentially harbor one or another type of DNMs.

Facioscapulohumeral muscular dystrophy (FSHD) region gene 1 (*FRG1*) (also other aliases) encodes a cytoplasmic protein relevant for muscular and vascular development. *FRG1* mapped 100 kb centromeric of the repeated units on chromosome 4q35 that contribute to the development of Facioscapulohumeral Muscular Dystrophy (FSHD, OMIM #158900)^46^. Although the molecular pathogenesis of FSHD remains unveiled, it is a muscular disease that shows comorbidity with several neurological symptoms such as epilepsy or ID^47^. It is believed that *FRG1* can act as a splicing regulator and it was recently reported that *Rbfox1* was downregulated when *FRG1* was overexpressed^48^. *RBFOX1* is involved in neuronal migration and synapse formation during corticogenesis and mutations in *RBFOX1* have been associated with ASD^49^. However, it should be taken into account that this gene is not constraint for truncating (pLI = 0) nor missense variants (Z = 0.18) and for that reason, this finding should be interpreted with caution. Thus, it is more probable that the dysregulation of target genes of FRG1 could be the cause of NDDs instead of its haploinsufficiency.

*KMT2C (lysine (K)-specific methyltransferase 2C)* encodes a methyltransferase that regulates gene transcription. *De novo* LoF mutations in *KMT2C* have been detected in individuals affected by Kleefstra syndrome 2 (OMIM #617768). Kleefstra syndrome is caused by haploinsufficiency of the euchromatin histone methyltransferase 1 (*EHMT1*) and characterized by delayed psychomotor development, ID, mild dysmorphic features and ASD. There is a molecular interplay between *KMT2C* and *EHMT1* both involved in the regulation of synaptic plasticity in the adult brain that could lead to the symptoms listed above^44^. In addition, mosaic mutations within *KMT2C* were previously identified when WES was carried out using novel pipelines that allow their detection^15^. These findings highlight the role of PZMs in some ASD individuals in genes previously identified as highest-confidence ASD risk genes.

*NFIA (Nuclear Factor I A)* encodes a member of the NF1 (nuclear factor 1) family of transcription factors determinants for the regulation of gliogenesis and other neuronal processes^50^. Although two DNMs in *NFIA* have been previously reported in patients with ID, there is no reliable evidence to consider it as a high-confidence ASD risk gene^51^. However, it is worth noting that an *NFIA*-related disorder caused by inherited or DNMs has been described. This syndrome is characterized by brain malformations and it can manifest or not renal urinary tract defects (OMIM #613735)^52^. The clinical presentation of the syndrome is highly variable and rarely all the described features are present in one individual^53^. To our knowledge, this is the first time that *NFIA* is pointed as a plausible candidate gene in ASD due to the analysis of PZMs that otherwise would have been dismissed.

The fact that different genes across the genome carry different types of DNMs (germinal and/or PZM), might be due to the fact that in some genes mutations are lethal in a germinal state but not in a mosaic state. This is the example of Rett’s Syndrome, in which *MECP*2 mutations in males are lethal and dominant in females, but in a few cases, mosaic mutations have been reported to be compatible with male viability^54^. However, taking into account our results, another reasonable hypothesis might be that the PZMs within some genes can cause different clinical manifestations that if they were germinal mutations. Accordingly, it was previously described that Kleefstra syndrome is caused by germinal LoF DNMs in *KMT2C* but PZMs located in the same gene cause milder phenotype presentation^55^. Thus, patients with less severe phenotypes and core ASD features are overrepresented in our cohort, allowing the detection of *KMT2C* association in the PZM analysis done with TADA, while patients carrying germinal DNMs in the same gene might have been excluded for presenting a syndromic form. For the same reason, *NFIA* could appear associated with ASD only when the mutations are mosaic.

From the biological point of view, the gene-set enrichment analysis done with both lists of genes harboring PZM and germinal mutations showed a significant enrichment for several gene-sets previously involved in ASD pathogenesis: FMRP targets, genes involved in chromatin organization and high-confidence genes from SFARI^6^. Curiously, only the list of PZM genes showed a significant enrichment for genes targeted by miR-137. miR-137 is a non-coding RNA well-known for its critical role during brain development^56^. The expression of miR-137 is crucial to maintain the balance between neuronal differentiation and proliferation. miR-137 is also involved in neuronal maturation, dendrite development and synaptogenesis^57–59^. In addition, common genetic variants within the gene that encodes miR-137 have been associated with ASD and schizophrenia^60,61^. In fact, the complete loss of miR-137 in mice is lethal, but a partial loss results in a phenotype which reproduces some of the core symptoms of ASD, such as repetitive behavior and impaired social functions^62^.

The GO enrichment analysis also points out different biological functions for those genes harboring germinal DNMs and PZMs^18,63^. Thus, GO terms mainly related to ion transport and modulation of the synaptic function were highlighted in germinal genes. However, PZM genes were enriched in GO terms mainly related to the negative regulation of gene expression.

Cell-type enrichment analysis and spatio-temporal expression analysis across brain regions and neurodevelopmental periods provide a novel insight into the differential role of PZM and germinal mutations. Germinal genes have shown a significant enrichment in both excitatory and inhibitory neurons while PZMs genes were mainly expressed in pyramidal neurons. In addition, it is worth to note that genes with germinal DNMs were expressed in both fetal and adult neurons across several brain areas (cortex, striatum, cerebellum and amygdala). However, the expression of genes carrying PZM was restricted to the mid-fetal cortex. Interestingly, the study performed by Lim et al. highlighted the amygdala as the main brain area in which PZMs were enriched for expression. However, this analysis only includes PZMs within critical exons, in comparison with the current study that includes all genes carrying non-synonymous PZMs^18^.

Taken together, these results suggest the existence of distinct mechanisms between PZM and germinal genes that could influence ASD susceptibility. Thus, impaired neuronal communication linked to mutations in germinal genes support the theory that excitatory/inhibitory imbalance contributes to ASD^21,64^. The fact that several brain areas are similarly affected during development might explain the clinical heterogeneity in ASD and the high comorbidity with other disorders such as epilepsy or ID in the presence of germinal variants. By contrast, although a replication in larger cohorts is needed, it has been suggested that individuals harboring PZMs might be less affected in terms of cognitive abilities. In these patients, some biological processes such as neurogenesis, neuronal migration and differentiation that occur within early developmental stages might be disrupted. However, this would only take place in some brain cells while the remaining cells will maintain a normal functioning. In agreement with this hypothesis, brains of children with ASD have shown patches of abnormal laminar organization that might be the result of the altered migration of just some cells to their target destination^65^.

Our results point out to a critical window during mid-fetal development in which the disruption of essential neurodevelopmental processes will take place. We demonstrated that ASD associated genes were expressed more often between mid-to-late fetal periods. In addition, although some genes harbor mutations in a mosaic state it is possible that the disruption of certain genes expressed during this crucial period would be enough to the later manifestation of ASD core symptoms.

## CONCLUSIONS

In conclusion, our analysis of germinal and PZM provide an additional insight into the role of PZMs in ASD etiology. It supports the previous evidence of the pathogenic role of PZM and their contribution to ASD risk. Moreover, our results suggest that PZMs might be implicated in different biological processes and neurodevelopmental stages than germinal mutations. In addition, these analyses revealed that ASD risk genes can differentially harbor PZMs or germinal mutations highlighting the importance of accurately detecting them in whole exome and genome studies as well as in clinical practice.

## LIST OF ABBREVIATIONS

AAF: alternate allele frequency
ADI-R: Autism Diagnostic Interview-Revised
ADOS: Autism Diagnostic Observation Schedule
ASD: Autism Spectrum Disorder
BEE: brain expressed enhancers
BF: bayesian factor
DNMs: de novo mutations
EWCE: Expression Weighted Cell type Enrichment
ExAC: Exome Aggregation Consortium
FDR: false discovery rate
FSHD: Facioscapulohumeral Muscular Dystrophy
GO: gene ontology
GQ: genotype quality
ID: intellectual disability
LoF: loss of function
NDD: neurodevelopmental disorder
OMIM: Online Mendelian Inheritance in Man
pSI: specifity index statistic
PZMs: postzygotic mutations
TADA: transmission and de novo association test
VCF: variant call format
WES: whole exome sequencing

## DECLARATIONS

## Acknowledgements

We would like to warmly thank the ASC (Autism Sequencing Consortium) (https://genome.emory.edu/ASC/) that has sequenced the 360 Spanish trios.

## Author’s contributions statement

JP, RM-R, MF-P and MP participated in the recruitment of samples. AA-G, CR-F, M-CC and JA have performed the analyses. AA-G wrote the paper. AC and CR-F critically revised the work and approved the final content. AA-G, CR-F, M-CC and AC participated in the design and coordination of this study.

## Ethics approval and consent to participate

The corresponding Research Ethics Committee of Galicia approved our study. All participants, parents or legal representatives provided written informed consent at enrollment and the study was conducted according to the declaration of Helsinki.

The ASC data employed in this study are already published (doi: 10.1038/nn.4598). The corresponding ethics committee has approved these genetic data.

## Consent for publication

Not applicable

## Availability of data and materials

All data generated during this study are included in this published article and its supplementary information files.

## REFERENCES

1. American Psychiatric Association. Diagnostic and Statistical Manual of Mental Disorders (Arlington, VA: American Psychiatric Publishing). (2013).

2. Lai, M.-C., Lombardo, M. V. & Baron-Cohen, S. Autism. Lancet Lond. Engl. 383, 896–910 (2014).

3. Sandin, S. et al. The Heritability of Autism Spectrum Disorder. JAMA 318, 1182–1184 (2017).

4. Gaugler, T. et al. Most genetic risk for autism resides with common variation. Nat. Genet. 46, 881–885 (2014).

5. Sanders, S. J. et al. Insights into Autism Spectrum Disorder Genomic Architecture and Biology from 71 Risk Loci. Neuron 87, 1215–1233 (2015).

6. De Rubeis, S. et al. Synaptic, transcriptional and chromatin genes disrupted in autism. Nature 515, 209–215 (2014).

7. Pinto, D. et al. Convergence of genes and cellular pathways dysregulated in autism spectrum disorders. Am. J. Hum. Genet. 94, 677–694 (2014).

8. Biesecker, L. G. & Spinner, N. B. A genomic view of mosaicism and human disease. Nat. Rev. Genet. 14, 307–320 (2013).

9. Bae, T. et al. Different mutational rates and mechanisms in human cells at pregastrulation and neurogenesis. Science 359, 550–555 (2018).

10. D’Gama, A. M. et al. Targeted DNA Sequencing from Autism Spectrum Disorder Brains Implicates Multiple Genetic Mechanisms. Neuron 88, 910–917 (2015).

11. Poduri, A., Evrony, G. D., Cai, X. & Walsh, C. A. Somatic mutation, genomic variation, and neurological disease. Science 341, 1237758 (2013).

12. D’Gama, A. M. & Walsh, C. A. Somatic mosaicism and neurodevelopmental disease. Nat. Neurosci. 21, 1504–1514 (2018).

13. Genovese, G. et al. Clonal hematopoiesis and blood-cancer risk inferred from blood DNA sequence. N. Engl. J. Med. 371, 2477–2487 (2014).

14. Pagnamenta, A. T. et al. Exome sequencing can detect pathogenic mosaic mutations present at low allele frequencies. J. Hum. Genet. 57, 70–72 (2012).

15. Freed, D. & Pevsner, J. The Contribution of Mosaic Variants to Autism Spectrum Disorder. PLoS Genet. 12, e1006245 (2016).

16. Dou, Y. et al. Postzygotic single-nucleotide mosaicisms contribute to the etiology of autism spectrum disorder and autistic traits and the origin of mutations. Hum. Mutat. 38, 1002–1013 (2017).

17. Krupp, D. R. et al. Exonic Mosaic Mutations Contribute Risk for Autism Spectrum Disorder. Am. J. Hum. Genet. 101, 369–390 (2017).

18. Lim, E. T. et al. Rates, distribution and implications of postzygotic mosaic mutations in autism spectrum disorder. Nat. Neurosci. 20, 1217–1224 (2017).

19. Lindhurst, M. J. et al. A mosaic activating mutation in AKT1 associated with the Proteus syndrome. N. Engl. J. Med. 365, 611–619 (2011).

20. Lee, J. H. et al. De novo somatic mutations in components of the PI3K-AKT3-mTOR pathway cause hemimegalencephaly. Nat. Genet. 44, 941–945 (2012).

21. Satterstrom, F. K. et al. Large-Scale Exome Sequencing Study Implicates Both Developmental and Functional Changes in the Neurobiology of Autism. Cell. 180, 568–584 (2020)

22. He, X. et al. Integrated model of de novo and inherited genetic variants yields greater power to identify risk genes. PLoS Genet. 9, e1003671 (2013).

23. Samocha, K. E. et al. A framework for the interpretation of de novo mutation in human disease. Nat. Genet. 46, 944–950 (2014).

24. Turner, S. qqman: an R package for visualizing GWAS results using Q-Q and manhattan plots. J. Open Source Softw. 3, 731 (2018).

25. Fromer, M. et al. De novo mutations in schizophrenia implicate synaptic networks. Nature 506, 179–184 (2014).

26. Darnell, J. C. et al. FMRP stalls ribosomal translocation on mRNAs linked to synaptic function and autism. Cell 146, 247–261 (2011).

27. Kirov, G. et al. De novo CNV analysis implicates specific abnormalities of postsynaptic signalling complexes in the pathogenesis of schizophrenia. Mol. Psychiatry 17, 142–153 (2012).

28. Georgi, B., Voight, B. F. & Bućan, M. From mouse to human: evolutionary genomics analysis of human orthologs of essential genes. PLoS Genet. 9, e1003484 (2013).

29. Cotney, J. et al. The autism-associated chromatin modifier CHD8 regulates other autism risk genes during human neurodevelopment. Nat. Commun. 6, 6404 (2015).

30. Lek, M. et al. Analysis of protein-coding genetic variation in 60,706 humans. Nature 536, 285–291 (2016).

31. Weyn-Vanhentenryck, S. M. et al. HITS-CLIP and integrative modeling define the Rbfox splicing-regulatory network linked to brain development and autism. Cell Rep. 6, 1139–1152 (2014).

32. Collins, A. L. et al. Transcriptional targets of the schizophrenia risk gene MIR137. Transl. Psychiatry 4, e404 (2014).

33. Wagnon, J. L. et al. CELF4 regulates translation and local abundance of a vast set of mRNAs, including genes associated with regulation of synaptic function. PLoS Genet. 8, e1003067 (2012).

34. Lin, M. et al. Allele-biased expression in differentiating human neurons: implications for neuropsychiatric disorders. PloS One 7, e44017 (2012).

35. Lelieveld, S. H. et al. Meta-analysis of 2,104 trios provides support for 10 new genes for intellectual disability. Nat. Neurosci. 19, 1194–1196 (2016).

36. Bartonicek, N. et al. Intergenic disease-associated regions are abundant in novel transcripts. Genome Biol. 18, 241 (2017).

37. Visel, A. et al. A high-resolution enhancer atlas of the developing telencephalon. Cell 152, 895–908 (2013).

38. Ching, A.-S. & Ahmad-Annuar, A. A Perspective on the Role of microRNA-128 Regulation in Mental and Behavioral Disorders. Front. Cell. Neurosci. 9, 465 (2015).

39. Merico, D., Isserlin, R., Stueker, O., Emili, A. & Bader, G. D. Enrichment map: a network-based method for gene-set enrichment visualization and interpretation. PloS One 5, e13984 (2010).

40. Dougherty, J. D., Schmidt, E. F., Nakajima, M. & Heintz, N. Analytical approaches to RNA profiling data for the identification of genes enriched in specific cells. Nucleic Acids Res. 38, 4218–4230 (2010).

41. Xu, X., Wells, A. B., O’Brien, D. R., Nehorai, A. & Dougherty, J. D. Cell type-specific expression analysis to identify putative cellular mechanisms for neurogenetic disorders. J. Neurosci. Off. J. Soc. Neurosci. 34, 1420–1431 (2014).

42. Jamuar, S. S. & Walsh, C. A. Somatic mutations in cerebral cortical malformations. N. Engl. J. Med. 371, 2038 (2014).

43. de Lange, I. M. et al. Mosaicism of de novo pathogenic SCN1A variants in epilepsy is a frequent phenomenon that correlates with variable phenotypes. Epilepsia 59, 690–703 (2018).

44. Koemans, T. S. et al. Functional convergence of histone methyltransferases EHMT1 and KMT2C involved in intellectual disability and autism spectrum disorder. PLoS Genet. 13, e1006864 (2017).

45. Schanze, I. et al. NFIB Haploinsufficiency Is Associated with Intellectual Disability and Macrocephaly. Am. J. Hum. Genet. 103, 752–768 (2018).

46. Hanel, M. L. et al. Facioscapulohumeral muscular dystrophy (FSHD) region gene 1 (FRG1) is a dynamic nuclear and sarcomeric protein. Differ. Res. Biol. Divers. 81, 107–118 (2011).

47. Saito, Y. et al. Facioscapulohumeral muscular dystrophy with severe mental retardation and epilepsy. Brain Dev. 29, 231–233 (2007).

48. Pistoni, M. et al. Rbfox1 downregulation and altered calpain 3 splicing by FRG1 in a mouse model of Facioscapulohumeral muscular dystrophy (FSHD). PLoS Genet. 9, e1003186 (2013).

49. Hamada, N. et al. Essential role of the nuclear isoform of RBFOX1, a candidate gene for autism spectrum disorders, in the brain development. Sci. Rep. 6, 30805 (2016).

50. Glasgow, S. M. et al. Glia-specific enhancers and chromatin structure regulate NFIA expression and glioma tumorigenesis. Nat. Neurosci. 20, 1520–1528 (2017).

51. Iossifov, I. et al. De novo gene disruptions in children on the autistic spectrum. Neuron 74, 285–299 (2012).

52. Lu, W. et al. NFIA haploinsufficiency is associated with a CNS malformation syndrome and urinary tract defects. PLoS Genet. 3, e80 (2007).

53. Revah-Politi, A. et al. Loss-of-function variants in NFIA provide further support that NFIA is a critical gene in 1p32-p31 deletion syndrome: A four patient series. Am. J. Med. Genet. A. 173, 3158–3164 (2017).

54. Pieras, J. I. et al. Somatic mosaicism for Y120X mutation in the MECP2 gene causes atypical Rett syndrome in a male. Brain Dev. 33, 608–611 (2011).

55. Kramer, J. M. et al. Epigenetic regulation of learning and memory by Drosophila EHMT/G9a. PLoS Biol. 9, e1000569 (2011).

56. Mahmoudi, E. & Cairns, M. J. MiR-137: an important player in neural development and neoplastic transformation. Mol. Psychiatry 22, 44–55 (2017).

57. Szulwach, K. E. et al. Cross talk between microRNA and epigenetic regulation in adult neurogenesis. J. Cell Biol. 189, 127–141 (2010).

58. Smrt, R. D. et al. MicroRNA miR-137 regulates neuronal maturation by targeting ubiquitin ligase mind bomb-1. Stem Cells Dayt. Ohio 28, 1060–1070 (2010).

59. He, E. et al. MIR137 schizophrenia-associated locus controls synaptic function by regulating synaptogenesis, synapse maturation and synaptic transmission. Hum. Mol. Genet. 27, 1879–1891 (2018).

60. Cross-Disorder Group of the Psychiatric Genomics Consortium. Identification of risk loci with shared effects on five major psychiatric disorders: a genome-wide analysis. Lancet Lond. Engl. 381, 1371–1379 (2013).

61. Ripke, S. et al. Genome-wide association analysis identifies 13 new risk loci for schizophrenia. Nat. Genet. 45, 1150–1159 (2013).

62. Cheng, Y. et al. Partial loss of psychiatric risk gene Mir137 in mice causes repetitive behavior and impairs sociability and learning via increased Pde10a. Nat. Neurosci. 21, 1689–1703 (2018).

63. Gompers, A. L. et al. Germline Chd8 haploinsufficiency alters brain development in mouse. Nat. Neurosci. 20, 1062–1073 (2017).

64. Nelson, S. B. & Valakh, V. Excitatory/Inhibitory Balance and Circuit Homeostasis in Autism Spectrum Disorders. Neuron 87, 684–698 (2015).

65. Stoner, R. et al. Patches of disorganization in the neocortex of children with autism. N. Engl. J. Med. 370, 1209–1219 (2014).

