## Additional file 4 for "Postzygotic and germinal *de novo* mutations in ASD: exploring their biological role"

Spain

^*^ These authors contributed equally to this work

| **Gene** | **qval** | **pval** | **Autism gene** | **SFARI gene** | **SFARI classification** | **Previous TADA; FDR <0.01** | **OMIM** | **OMIM Number** |
| --- | --- | --- | --- | --- | --- | --- | --- | --- |
| SCN2A | 5.04 x 10^-12^ | 4.13 x 10^-8^ | yes | yes | 1 | yes | Epileptic encephalopathy, early infantile, 11; Seizures, benign familial infantile, 3 | #613721,# 607745 |
| CHD8 | 2.40 x 10^-5^ | 4.13 x 10^-8^ | yes | yes | 1s | yes | {Autism, susceptibility to, 18} | #615032 |
| ARID1B | 5.36 x 10^-5^ | 4.13 x 10^-8^ | yes | yes | 1s | yes | Coffin-siris syndrome 1 | #135900 |
| SLC6A1 | 0.00014991 | 2.48 x 10^-7^ | yes | yes | 2s | yes | Myoclonic-atonic epilepsy | #616421 |
| SYNGAP1 | 0.000508219 | 6.61 x 10^-7^ | yes | yes | 1s | yes | Mental retardation, autosomal dominant 5 | #612621 |
| KDM5B | 0.000841602 | 8.26 x 10^-7^ | yes | yes | 2s | no | Mental retardation, autosomal recesyesve 65 | #618109 |
| KMT5B | 0.001898533 | 5.37 x 10^-6^ | yes | yes | 1S | yes | Mental retardation, autosomal dominant 51 | #617788 |
| TRIP12 | 0.0027703 | 5.79 x 10^-6^ | yes | yes | 1S | no | Mental retardation, autosomal dominant 49 | #617752 |
| PTEN | 0.004131864 | 1.14 x 10^-5^ | yes | yes | 1S | yes | Cowden syndrome 1;Lhermitte-Duclos syndrome;Macrocephaly/autism syndrome;Prostate cancer, somatic;{Glioma susceptibility 2};{Meningioma} | #158350; #158350; #605309; #176807;# 613028; #607174 |
| KATNAL2 | 0.008224845 | 5.01 x 10^-5^ | yes | yes | 1 | yes | NA | NA |
| NRXN1 | 0.01194415 | 5.79 x 10^-5^ | yes | yes | 2 | no | Pitt-Hopkins-like syndrome 2; {Schizophrenia, susceptibility to, 17} | #614325; #614332 |
| CREBBP | 0.015098004 | 5.92 x 10^-5^ | no | yes | 5 | no | Menke-Hennekam syndrome 1;Rubinstein-Taybi syndrome 1 | #618332;#180849 |
| CELF4 | 0.017826919 | 6.09 x 10^-5^ | no | yes | 3 | no | NA | NA |
| STXBP1 | 0.020341344 | 6.48 x 10^-5^ | no | yes | 3s | NO | Epileptic encephalopathy, early infantile, 4 | #612164 |
| DYRK1A | 0.0227959 | 7.09 x 10^-5^ | yes | yes | 1s | yes | Mental retardation, autosomal dominant 7 | #614104 |
| CHD2 | 0.025896407 | 0.00010719 | yes | yes | 1s | no | Epileptic encephalopathy, childhood-onset | #615369 |
| ANK2 | 0.029572774 | 0.000136694 | yes | yes | 1 | yes | Cardiac arrhythmia, ankyrin-B-related; Long QT syndrome 4 | #600919; #600919 |
| WDFY3 | 0.033145437 | 0.000147769 | yes | yes | 2 | no | ?Microcephaly 18, primary, autosomal dominant | #617520 |
| UNC80 | 0.037280204 | 0.00018595 | no | yes | 4 | no | Hypotonia, infantile, with psychomotor retardation and characteristic facies 2 | #616801 |
| CLASP1 | 0.041141977 | 0.000192149 | no | yes | 3 |  | NA | NA |
| TMEM39B | 0.045179962 | 0.000226612 | no | no | NA | no | NA | NA |
| PRKAR1B | 0.049032281 | 0.000240909 | no | no | NA | no | NA | NA |
| USP45 | 0.053777544 | 0.000349174 | no | yes | 3 | no | NA | NA |
| NUAK1 | 0.058346344 | 0.000365372 | no | yes | 3 | no | NA | NA |
| NAA15 | 0.062733828 | 0.000387769 | yes | yes | 1s | yes | Mental retardation, autosomal dominant 50 | #617787 |
| FOXP1 | 0.066799738 | 0.000390331 | yes | yes | 1s | no | Mental retardation with language impairment and with or without autistic features | #613670 |
| ZC3H11A | 0.07061159 | 0.00039876 | no | no | NA | no | NA | NA |
| DPP3 | 0.074752544 | 0.000470909 | no | no | NA | no | NA | NA |
| PRKDC | 0.079128953 | 0.00053124 | no | yes | 4 | no | Immunodeficiency 26, with or without neurologic abnormalities | #615966 |
| ATP1A1 | 0.083428257 | 0.000549008 | no | yes | 4s | no | Charcot-Marie-Tooth disease, axonal, type 2DD; Hypomagnesemia, seizures, and mental retardation 2 | 618036; 618314 |
| LRP5 | 0.087492397 | 0.000553223 | no | no | NA | no | Exudative vitreoretinopathy 4; Hyperostoyess, endosteal; Osteopetroyess, autosomal dominant 1; Osteoporoyess-pseudoglioma syndrome; Osteoscleroyess; Polycystic liver disease 4 with or without kidney cysts; van Buchem disease, type 2; [Bone mineral denyesty variability 1]; {Osteoporoyess} 166710 AD 3 | #601813; #144750; #607634; #259770; #144750; #617875; #607636; #601884; #166710 |
| SLC12A3 | 0.091716102 | 0.000608099 | no | no | NA | no | Gitelman syndrome | #263800 |
| FBXO18 | 0.095909862 | 0.000644628 | no | no | NA | no | NA | NA |
| PTK7 | 0.099893315 | 0.000651157 | no | yes | 3 | yes | NA | NA |

**Supplementary table 20. Description of germinal targeted genes obtained with TADA in the combined cohort.** Germinal targeted genes with FDR values <0.1 are shown. The following information is included: Gene already recognized as a true ASD gene, Gene included in the SFARI dataset, Genes previously found in TADA analysis and OMIM disease associated to the gene.

| **Gene** | **qval** | **pval** | **Autism gene** | **Sfari** | **SFARI classification** | **Previous TADA; FDR <0.01** | **OMIM** | **OMIM Number** |
| --- | --- | --- | --- | --- | --- | --- | --- | --- |
| FRG1 | 0.035045881 | 4.14 x 10^-5^ | no | no | NA | no | Facioscapulohumeral muscular dystrophy 1 | #158900 |
| KMT2C | 0.069130706 | 0.000182873 | yes | yes | s2 | yes | Kleefstra syndrome 2 | #617768 |
| NFIA | 0.091745554 | 0.000279834 | no | yes | 4 | no | Brain malformations with or without urinary tract defects | #613735 |
| SMARCA4 | 0.118024039 | 0.000517127 | no | yes | 3 | no | Coffin loris | #614609 |
| PRKDC | 0.13475291 | 0.000545856 | no | yes | 4 | no | Immunodeficiency 26, with or without neurologic abnormalities | #615966 |
| KLF16 | 0.149512294 | 0.00064116 | no | yes | 4 | no | NA | NA |
| GRIN2B | 0.169042489 | 0.000949171 | yes | yes | 1 | yes | Epileptic encephalopathy, early infantile, 27; Mental retardation, autosomal dominant 6 | #616139, #613970 |
| MAP2K3 | 0.184127855 | 0.000977624 | no | no | NA | NA | NA | NA |
| HNRNPU | 0.209192662 | 0.001851381 | no | yes | 4 | no | Epileptic encephalopathy, early infantile, 54 | #617391 |
| POTEB2 | 0.23133466 | 0.002020718 | no | no | NA | NA | NA | NA |
| RNPC3 | 0.250159757 | 0.002077072 | no | no | NA | NA | ?Growth hormone deficiency, isolated, type V | #618160 |
| FAM177A1 | 0.26825174 | 0.002417127 | no | no | NA | NA | NA | NA |
| CALML6 | 0.28374479 | 0.002460773 | no | no | NA | NA | NA | NA |
| CMPK2 | 0.297509845 | 0.002585083 | no | no | NA | NA | NA | NA |

**Supplementary table 21. Description of PZMs targeted genes obtained with TADA in the combined cohort.** PZMs targeted genes with FDR values <0.3 are shown. The following information is included: Gene already recognized as a true ASD gene, Gene included in the SFARI dataset, Genes previously found in TADA analysis and OMIM disease associated to the gene.

| Term | P-value | Adjusted P-value | Genes |
| --- | --- | --- | --- |
| sodium ion transport (GO:0006814) | 2.043 x 10^-6^ | 0.005180034 | SLC12A3;ATP1A4;SLC4A11;ANO6;PKD2L1;SLC5A1;SLC4A4;SLC8A2;SCNN1G;SCNN1D;SCN11A;SCN7A;SCN4A;SLC17A1;SCN2A;SCN3A;HCN2;SLC17A3;SCN1A |
| beta-catenin-TCF complex assembly (GO:1904837) | 2.997 x 10^-6^ | 0.005180034 | TLE4;TCF7L2;CREBBP;TLE2;TLE1;TRRAP;LEF1;TERT;RBBP5;RUVBL1;LEO1;CTNNB1;EP300 |
| regulation of transcription from RNA polymerase II promoter (GO:0006357) | 5.112 x 10^-6^ | 0.005180034 | ATF2;MYT1L;NOC2L;TIAL1;TBK1;RPS6KA1;MYB;SMARCC1;MEF2C;SMARCC2;PRKCB;RFX2;DICER1;MED4;RFX8;TCERG1L;RFX7;ZSCAN21;PRKD2;PRKD1;ZFPM2;HOXB7;SRCAP;DHX9;PRKDC;CTR9;GATA4;RTF1;ZBTB5;MED12L;TP53BP1;ZNF268;STAT5A;CREBBP;BCL11B;ZNF382;SMARCA5;NR2F1;NFATC1;SMARCA2;NFATC4;MED13L;POU6F1;SETX;AGO1;CDK13;RCOR2;GTF2H2C;TCF20;CTCF;BRCA1;PCSK6;GLI3;IKBKB;MECOM;HEY1;SUFU;CIC;TEAD1;ZNF366;HTATIP2;NCOA1;FOXD3;ZHX3;NCOA6;FSHB;TET2;TET1;PPRC1;SUPT3H;RGMA;NCOR2;KAT2A;MED21;MTF2;RARA;ARHGEF2;TLR4;JDP2;TMPRSS6;SATB1;DOT1L;TRAK1;EGFR;HIRA;CUX2;CUX1;NSD1;LEO1;MEF2D;BRD4;MBD4;ZNF462;NR1H2;STAT2;DAB2IP;ZNF76;SKI;PER1;FAM200B;PSMC5;NFIA;KLHL6;NFIB;ZNF219;CTNNB1;PKN1;MEGF8;CRLF3;KIAA1958;CRTC3;CD81;RORC;RORA;RORB;TTF1;FGF2;RUVBL1;SLC39A5;EP300;ZNF568;KDM2B;HGF;ZBTB38;FOXP2;FOXP1;PHIP;DDX5;NOTCH1;AKNA;TNKS;NPAS3;RAI1;DEAF1;NLRP3;MAP2K5;BPTF;ZBTB17;ZFHX3;SMAD4;TFAP2C;CBX4;WFS1;SMURF1;SMAD6;BMP3;AHI1;SP2;SP3;CAPRIN2;MDM2;SP7;KDM5B;ZC3H4;CHD8;ONECUT1;CHD5;CHD4;CHD2;PKD1;FLCN;SIN3B;SIN3A;DACT1;KDM6B;MYOCD;RBM14;DDX58;MKL1;VEZF1;ETV1;ETV2;KLF17;ETV6;PPM1A;SFPQ;THRAP3;CRY2;CDH13;MET;MCPH1;LEF1;CEBPG;TAF9;DDX20;LRP5;CXXC5;HDAC9;BEND3;MAPK3;SPEN;SMARCE1;TCF7L2;MACC1;ATP2B4;ASXL3;TRIM37 |
| sodium ion transmembrane transport (GO:0035725) | 5.358 x 10^-6^ | 0.005180034 | SLC12A3;ANO6;PKD2L1;SLC8A2;SCNN1G;SCN11A;SCN7A;SCN4A;SLC17A1;SCN2A;SCN3A;HCN2;SLC17A3;SCN1A |
| positive regulation of excitatory postsynaptic potential (GO:2000463) | 6.195 x 10^-6^ | 0.005180034 | NLGN1;RELN;CUX2;NLGN2;CHRNA7;NRXN1;PTEN;PTK2B;SHANK3;SHANK1 |
| ion transport (GO:0006811) | 7.397 x 10^-6^ | 0.005180034 | GABRB3;RYR1;GABRB2;RYR2;ATP8A1;ATP2A1;PKD2L1;SLC4A2;RYR3;SLC8A2;UNC80;PANX2;PIEZO2;CNGA3;CHRNB1;SLC15A2;ANO9;ATP11B;ANO6;ANO5;GABRG2;GABRG1;SCNN1G;SLC5A8;SCNN1D;CNGB3;CNGB1;ATP6V1A;SLC24A3;SLC26A1;CHRNA7;ATP1A4;CLCNKB;CLCNKA;ATP1A3;ATP10A;ATP1A1;CLCA2;SLC17A1;CLCA4;SLC17A3;SLC12A3;GABRA1;SLC12A4;ATP8B2;ATP8B1;ATP2B4;CLCN7;ATP4A;P2RX5;P2RX1;SLC26A7;CHRFAM7A |
| calcium ion transport (GO:0006816) | 1.008 x 10^-5^ | 0.006048506 | RYR1;RYR2;SLC24A3;CATSPER4;CHRNA7;CAMK2A;ATP2A1;CACNA1D;CACNA1E;PKD1;RYR3;CACNA1H;SLC8A2;TRPM1;CYP27B1;CACNA1I;CCL5;CDH23;ITGAV;TRPM7;CACNA1S;TRPM6;TRPM4;TRPC4;CACNA2D1;ATP2B4;ANK2;ANO6 |
| modulation of chemical synaptic transmission (GO:0050804) | 1.196 x 10^-5^ | 0.006242463 | USP46;GRIA1;GRIA2;NLGN1;GRID2;MEF2C;NLGN2;SYT1;CHRNA7;GRIK5;CAMK2A;GRIK3;PTEN;GRIK4;GRIK1;BTBD9;SNCAIP;KBTBD4;PSMC5;RELN;GRM7;CNTNAP4 |
| phosphorylation (GO:0016310) | 1.337 x 10^-5^ | 0.006242463 | GSK3B;PANK2;MAST2;STK19;MYLK3;IKBKB;TBK1;GNPTAB;RPS6KA1;CDK20;TLK1;NEK1;NEK2;MAP4K1;MAP2K1;MORC3;CSNK2A1;PRKCB;MINK1;DYRK1A;LMTK2;PRKCA;PRPF4B;CSNK1E;PASK;ZAP70;ERN2;PRKAR1B;RARA;PRKD2;BIRC6;TOP1;TNIK;UQCRC2;PRKD1;BRSK1;LTK;BRSK2;PRKDC;CAMK2A;NUAK1;STK36;ABL1;PTK2B;MARK2;MAPK3;TEFM;SRPK2;DMPK;INSR;GLYCTK;LIMK1;EIF2AK2;BRAF;CDC42BPB;CDC42BPA;MERTK;GNPTG;PINK1;TAOK3;TAOK1;NEK10;PKN1;CDK13;MAP3K11 |
| modulation of excitatory postsynaptic potential (GO:0098815) | 1.581 x 10^-5^ | 0.006642747 | NLGN1;MTMR2;RELN;CUX2;NLGN2;CHRNA7;NRXN1;PTEN;PTK2B;SHANK3;SHANK1 |

**Supplementary table 22. Top 10 biological processes enriched in the germinal set.** Gene ontology enrichment analysis was done in germinal targeted genes from the combined cohort. Top 10 biological processes are shown.

| Term | P-value | Adjusted P-value | Genes |
| --- | --- | --- | --- |
| negative regulation of gene expression (GO:0010629) | 2.78671E-05 | 0.07390359 | NOTCH2;RB1;L3MBTL1;ZNF253;NDUFA13;HDAC1;CHD8;ZBTB21;BRCA1;GIGYF2;BCLAF1;NIPBL;DEAF1;PRDM16;ENC1;KDR;SOX9;CIC;DIS3L2;SUZ12;TCF7L2;MEF2C;FBXW11;ZBTB38;MBD2;HMGA1;PML;SMARCA4;EID1;TFAP4;PC;AEBP2;TRAF6;SP3;MTF2;RARA;ATF5;ZFPM2;TAF1 |
| negative regulation of cellular macromolecule biosynthetic process (GO:2000113) | 7.71495E-05 | 0.092248288 | RB1;L3MBTL1;ZNF253;NDUFA13;HDAC1;CHD8;ZBTB21;BRCA1;GIGYF2;BCLAF1;NIPBL;DEAF1;PRDM16;ENC1;SOX9;CIC;TCF7L2;FBXW11;ZBTB38;MBD2;HMGA1;PASK;PML;GTPBP4;SMARCA4;EID1;TFAP4;TRAF6;SP3;RARA;ATF5;ZFPM2;ATR |
| negative regulation of centriole replication (GO:0046600) | 0.000104353 | 0.092248288 | KAT2B;CHMP2A;BRCA1;MDM1 |
| regulation of megakaryocyte differentiation (GO:0045652) | 0.000144905 | 0.096071978 | L3MBTL1;KAT2B;KMT2D;MEF2C;HDAC1;KMT2C;THBS1;TNRC6B |
| response to calcium ion (GO:0051592) | 0.000187599 | 0.099502294 | MEF2C;CASR;AHCYL1;RASA4;PDCD6;HPCA;ITPR3;THBS1;TTN;SLC25A13 |
| negative regulation of nucleic acid-templated transcription (GO:1903507) | 0.000377487 | 0.135689506 | RB1;L3MBTL1;ZNF253;NDUFA13;HDAC1;CHD8;ZBTB21;BRCA1;BCLAF1;NIPBL;DEAF1;PRDM16;SOX9;CIC;TCF7L2;FBXW11;ZBTB38;MBD2;HMGA1;PML;SMARCA4;EID1;TFAP4;TRAF6;SP3;RARA;ATF5;ZFPM2 |
| protein phosphorylation (GO:0006468) | 0.000425767 | 0.135689506 | PRKDC;MAST1;WNK4;PKD1;EGFR;KDR;PIM1;RICTOR;EPHB2;ROS1;MAP4K3;PRKCG;CDK19;CAMK1D;CDC42BPG;HUNK;CDC7;GTF2H1;PRPF4B;CDC42BPB;PASK;DCLK1;LATS1;RARA;BIRC6;CDK13;TLR2;ATR;TAF1 |
| negative regulation of cell migration (GO:0030336) | 0.000436857 | 0.135689506 | IFITM1;PTPRR;ADAM15;NAV3;KRT16;DAG1;PTEN;EPPK1;SRGAP2;ARID2;GTPBP4;PTPRG |
| MAPK cascade (GO:0000165) | 0.000510044 | 0.135689506 | PLVAP;MEF2C;PSPN;FBXW11;CSF2RB;RASAL1;GRIN2B;EGFR;PSMD8;PSMB4;PPP5C;SYNGAP1;RASA4;TPR;IL3RA;KDR;PSME2;SOX9;SCG2;MAP4K3 |
| central nervous system development (GO:0007417) | 0.00051165 | 0.135689506 | PSPN;NCOA6;CHD8;GSTP1;PTEN;TBR1;CELSR1;GRIN2B;PKD1;DCLK1;ACAN;NIPBL;ALDH5A1;RARA;PLXNB2;CIC;MDGA1 |

**Supplementary table 23. Top 10 biological processes enriched in the PZMs set.** Gene ontology enrichment analysis was done in PZMs targeted genes from the combined cohort. Top 10 biological processes are shown.

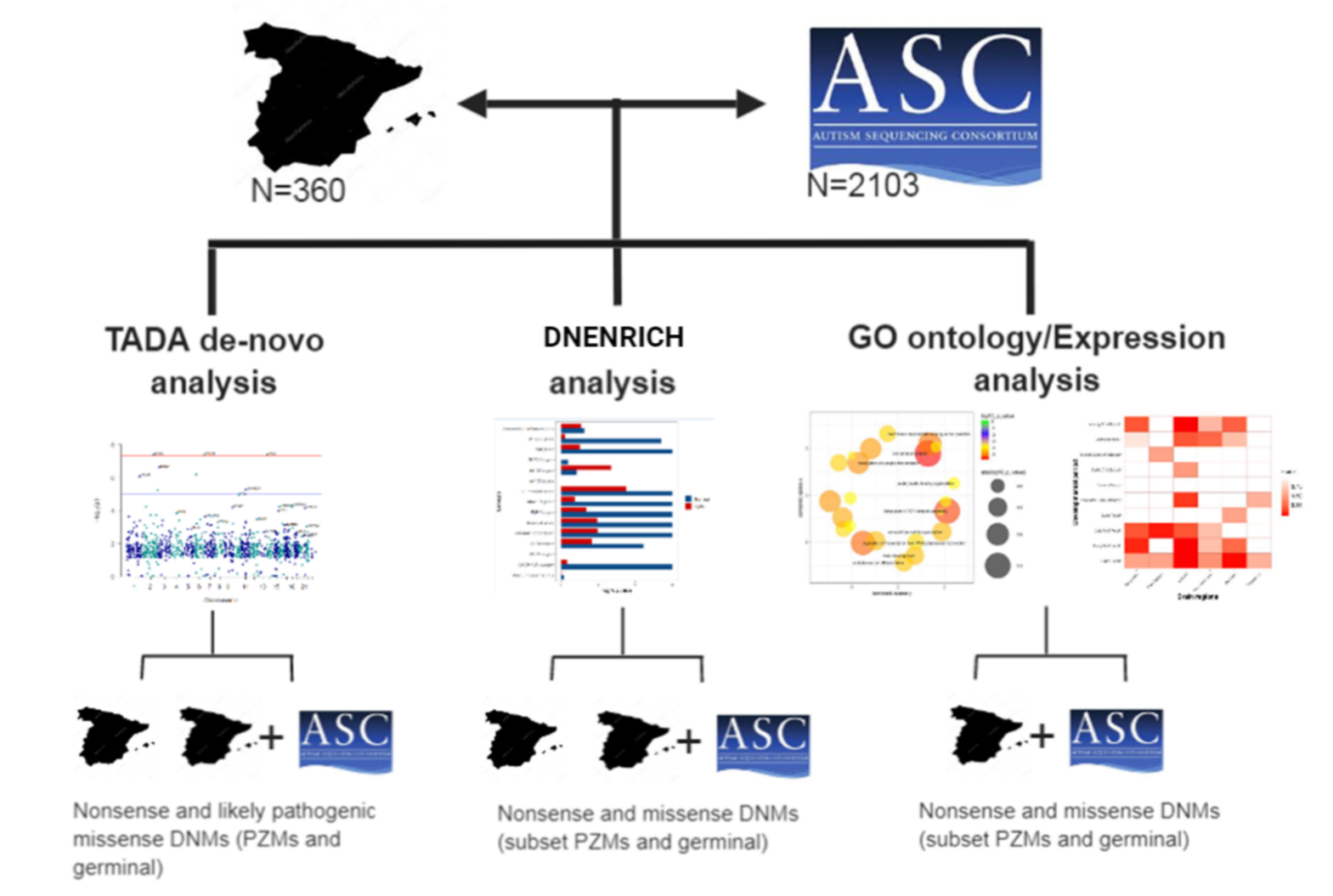
**Supplementary figure 1**. **Graphical view of the analysis developed in the Spanish and combined cohort.** TADA de-novo analysis was done in genes carrying germinal and PZMs DNMs in both cohorts considering nonsense and likely pathogenic missense variants. DNENRICH analysis was also done in genes carrying germinal DNMs and PZMs in both cohorts considering nonsense and missense variants. Go ontology enrichment analysis and expression analysis was done in genes carrying germinal DNMs and PZMs in the combined cohort, considering nonsense and missense variants.
